# Soft and stretchable organic bioelectronics for continuous intra-operative neurophysiological monitoring during microsurgery

**DOI:** 10.1101/2022.11.28.518170

**Authors:** Wenjianlong Zhou, Yuanwen Jiang, Qin Xu, Liangpeng Chen, Hui Qiao, Yixuan Wang, Jiancheng Lai, Donglai Zhong, Yuan Zhang, Weining Li, Yanru Du, Xuecheng Wang, Jiaxin Lei, Gehong Dong, Xiudong Guan, Shunchang Ma, Peng Kang, Linhao Yuan, Milin Zhang, Jeffrey B.-H. Tok, Deling Li, Zhenan Bao, Wang Jia

## Abstract

Continuous intra-operative neurophysiological monitoring (CINM) that provides precise mapping of neural anatomy through the entire microsurgery process is essential to preserve the structural and functional integrity of the nerve. However, bulky and rigid electrodes used currently in clinics are usually unable to reliably maintain continuous and stable contacts with the vulnerable and complex nerve networks, thus often resulting in detrimental post-operative complications, such as hemiplegia and sensory disturbances. Here, we describe a biomechanically compatible, suture-free, and individually reconfigurable CINM based on soft and stretchable organic electronic materials. Due to both low impedance and modulus of our conducting polymer electrodes, we achieved *for the first time* continuous recording of near-field action potential with high signal-to-noise ratio and minimal invasiveness during microsurgeries. Utilizing this unprecedented CINM modality, in conjunction with localized neurostimulation, we further demonstrated our approach in enabling optimal post-operative prognosis in preclinical animal models by preserving normal neural functions after a variety of tumor resection surgeries.

## Introduction

Nervous system neoplasm is the most common type of tumor affecting our nervous system with high associated morbidity and mortality^1,2^. In 2021, ~85,000 patients were diagnosed with primary malignant brain and other central nervous system neoplasms in the United States, and ~22% of the patients died from the disease^3^. Currently, the most effective treatment method of neurological neoplasm is through surgical excision of the tumor^4–7^. However, since tumoral lesions are typically associated with significant disruptions of the delicate nerve structures, even with meticulous surgical operations, post-operative complications such as nerve paralysis, amyotrophy, and sensory disorders remain serious issues^8,9^. To preserve the structural and functional integrity of the nerves after tumor resection, continuous intra-operative neurophysiological monitoring (CINM) throughout the entire microsurgery process is essential^5,10–12^. To date, CINM techniques used in clinics mostly rely on far-field potentials, which include electromyogram (EMG), electroencephalography (EEG), and brainstem auditory evoked potentials (BAEP)^10–13^. However, these far-field signals are typically weak, requiring long-time acquisition and summation to achieve discernable signal-to-noise ratios. As a result, far-field potential recording itself is insufficient to provide high-quality physiological monitoring in real time^14–16^. In addition, many tumor-bearing patients already have considerable pre-operative neurological symptoms, which further lower the source neural signals. Moreover, the far-field potential waveform is also sensitive to environmental variations, e.g., hypothermia or hypocarbia, making it extra challenging for traditional neurological monitoring approaches^17^.

On the other hand, near-field potentials evoked directly from the target nerves would provide instantaneous feedback with much higher amplitudes (typically >20 times higher than far-field signals)^14,18–21^. Early detection of abnormal signals enables early management to prevent irreversible modifications to the neural pathway. In principle, continuous monitoring of near-field potentials during neurosurgery will be the ideal method to assess neurological functions and improve prognosis (**Fig. 1a**). In practice, it is however highly challenging to have stable and high-quality monitoring throughout the entire surgery. This is because the highly complicated and vulnerable nerve networks require the neural interfacing electrodes to have multiple channels, high mechanical compliance with robust contacts at the same time. For clinically used ball-point electrodes made of stainless steel, due to their high stiffness and bulky size, they have limited mechanical compatibility with the delicate tissues, which may cause irreparable damage during probe movement. In addition, these hand-held electrodes could only be used intermittently during the surgery, making them inappropriate for continuous monitoring (**Fig. 1b**). Although new developments based on thin metal wires have been attempted^18,19^, none of the existing probes can maintain stable and reliable recording during routine neurosurgical manipulations, as they inevitably are subjected to tugging, displacing, and bending motions. To address the limitations of current CINM devices^14,18,19,22^, we developed a biomechanically compatible, suture-free, and individually reconfigurable CINM based on soft and stretchable organic bioelectronic devices (Fig. S1). Compared to rigid metal electrodes, conducting polymer electrodes with low modulus can ‘physically conform’ to the soft brain tissues and therefore significantly decrease the mechanical mismatch and improve signal quality^23^. In addition, conducting polymers with their electronic-ionic dual conduction as well as large volumetric capacitance can further reduce the interfacial impedance^24^. To demonstrate the efficacy of our device for CINM, we focused on the schwannoma, which is the most common of all nerve sheath tumors^25,26^. Neural preservation is particularly imperative in the management of schwannoma. Within the schwannoma family, the most prevalent type is vestibular schwannomas (VS), which accounts for ~60% of total cases^4,27–29^. Without CINM of near field signals evoked from the auditory nerves, i.e., cochlear nerve action potentials (CNAP), ~70% of the patients after the surgery would suffer from ipsilateral sensorineural hypoacusis^30,31^. In this work, we describe a new surgical scheme to implant the soft and stretchable conducting polymer electrodes. Using this approach, we achieved *for the first time* stable CNAP recording with minimal invasiveness during the entire VS resection, yielding optimal postoperative prognosis in preclinical animal models (**Fig. 1c**). Utilizing the unprecedented CINM modality in conjunction with localized neurostimulation, we further demonstrated our capability in restoring normal neural functions after a variety of neurosurgeries.

**Fig. 1.**
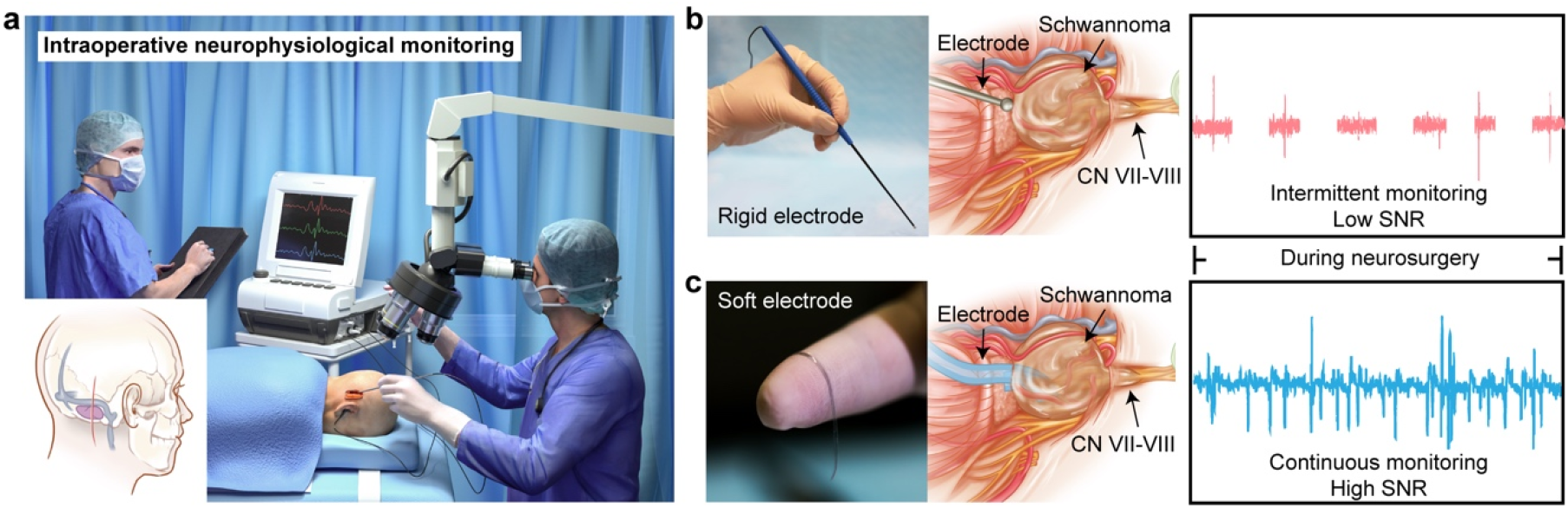
Soft PEDOT electrodes for CINM. **a,** Schematics showing CINM during neurosurgery. **b,** Images and schematics of INM using conventional rigid ball-point electrodes. The bulky electrode could only be placed on the cisternal portion of the cochlear nerve proximal to the cerebellopontine angle tumor. In addition, these hand-held electrodes could only be used intermittently during the surgery, making them not feasible for continuous monitoring. **c,** Images and schematics of CINM using soft PEDOT electrodes. Stable enclosure and CINM were achieved by wrapping a PEDOT electrode around the facial-acoustic nerve complex. CINM: Continuous intra-operative neurophysiological monitoring. SNR: Signal-to-noise ratio.

## Translatable surgical procedures

In this work, we used poly(3,4-ethylenedioxythiophene):polystyrene sulfonate (PEDOT:PSS) as the electrode material because it possesses the best electrical properties among all conducting polymers^32^, as well as its excellent biocompatibility^33^. By blending PEDOT:PSS with a previously designed crosslinkable supramolecular additive, we prepared paper-thin rail electrodes with both high electrical conductivity and mechanical stretchability^34^. Due to the low modulus of the device, we can simply wrap it around the nerve to create a soft enclosure, such that the PEDOT electrode can naturally fit the nerve complex with tight tissue-electrode contact (**Fig. 2a**). To demonstrate the potential of the soft PEDOT electrodes for clinical translation, we showed that the device can be easily implanted in a human cadaver skull via a retrosigmoid approach to expose the facial-acoustic nerve complex (**Fig. 2b**, Fig. S2). Retrosigmoid craniotomy is routinely performed during VS microsurgical resection because it provides a wide view of the cerebellopontine angle (CPA), which covers the majority of the cranial nerves from the fourth to the eleventh (Fig. S3a)^35^.

**Fig. 2.**
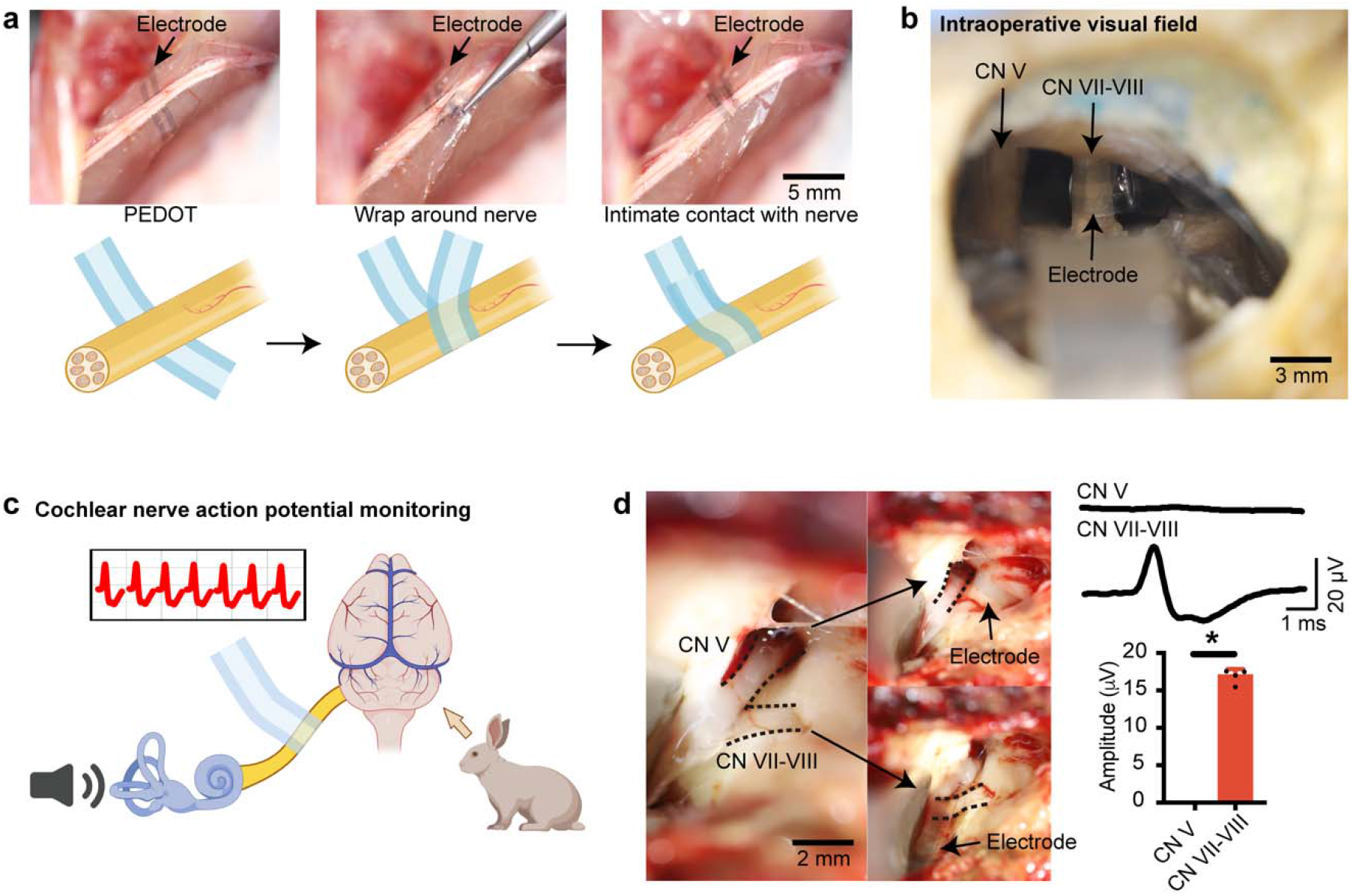
Soft PEDOT electrodes for conformable neural interfaces. **a,** Schematics and images of the implantation process for soft PEDOT electrode. Stable enclosure was achieved by wrapping a soft PEDOT electrode around the sciatic nerve and pressing gently without noticeable nerve tugging. **b,** Photograph of a soft PEDOT electrode wrapping around the facial-acoustic nerve complex in a human cadaver skull using a retrosigmoid approach. **c,** Schematics showing intra-operative CNAP monitoring during VS surgery in a rabbit model. **d,** Isolation of the facial-acoustic nerve complex from the cranial nerves. Two unidentified nerves were exposed after retracting the cerebellum of an anesthetized rabbit. Soft PEDOT electrodes were wrapped around each visible nerve for facial-acoustic nerve complex identification (*n* = 4, *P* = 0.029). CNAP values were measured from the PEDOT device. All error bars denote s.d. \**P* < 0.05; \*\**P* < 0.01; \*\*\**P* < 0.001; unpaired, two-tailed Student’s t-test was used for d. CNAP: Cochlear nerve action potentials; VS: Vestibular schwannomas.

Besides establishing the desired neural interfaces, another key advantage of our PEDOT electrode (versus what is currently used in clinics) is that we could easily multiplex the number of channels to quickly identify the nerve of interest during the surgical procedure^34^. Typically, for patients with brain tumors, because of the unrestricted tumor outgrowth, their cranial nerves are usually severely distorted with highly disrupted arrangements (**Supplementary Fig. 3b-c**). As a result, it remains a challenging task for neurosurgeons to distinguish the target nerve, e.g., the cochlear nerve in the VS case. By placing our fabricated PEDOT electrode array around multiple nerves, we could effectively send stimulus through individual channels of the device and hence readily identify the facial-acoustic nerve complex in a rabbit model (**Fig. 2c-d,** Figs. S4-5).

## Stable and continuous intra-operative auditory monitoring

To evaluate the device’s performance in monitoring neurophysiological signals during craniotomy, we performed two separate measurements in the rabbit model, i.e., BAEP (far-field) and CNAP (near-field). BAEP is the current gold standard for CINM during VS resection surgery, in which it is recorded on the scalp following sound stimulation of the ear. CNAP, on the other hand, is recorded directly from the auditory nerve. Prior to the microsurgery, by sweeping the stimulus sound frequencies to establish the baseline signals, we observed that 16 kHz inputs could evoke the most prominent responses with stable waveforms across all decibels. This acoustic property was then used for the entire experiments (Figs. S6-S7).

Using commercial needle electrodes to monitor BAEP and our PEDOT device for CNAP, we first showed that both BAEP and CNAP can be reliably recorded before and after the craniotomy (**Fig. 3a-e**). The marginal differences between signal amplitudes and latencies from pre- and post-operative recordings confirmed minimal damage on hearing during the microsurgery (Fig. S8). However, when comparing both CNAP and BAEP signals, CNAP had an average amplitude of ~15 μV whereas BAEP was only ~300 nV due to its far-field nature (**Fig. 3f**). Due to the >50-fold differences in the signal amplitudes, the required acquisition durations for each type of data were also drastically different. For example, it took at least 5 s to collect a set of BAEP waveforms because of the multi-time repeats needed for averaging (Video S1). On the other hand, CNAP signals can be acquired in <500 ms, which was one order of magnitude faster than BAEP (Video S2). The improved temporal resolution was essential to capture any possible damages caused by surgical maneuvers and to minimize the risk of hearing loss. To mimic the ‘extreme’ scenarios during surgical procedures, we intentionally tugged the nerve using a stripper and examined the dynamic changes of BAEP or CNAP. Compared to the slow response (10.5 s) and long recovery latency (10.6 s) of BAEP signals, CNAP recorded using PEDOT showed orders of magnitude improvements of 0.52 s (response latency) and 0.58 s (recovery latency), respectively (**Fig. 3g-j**).

**Fig. 3.**
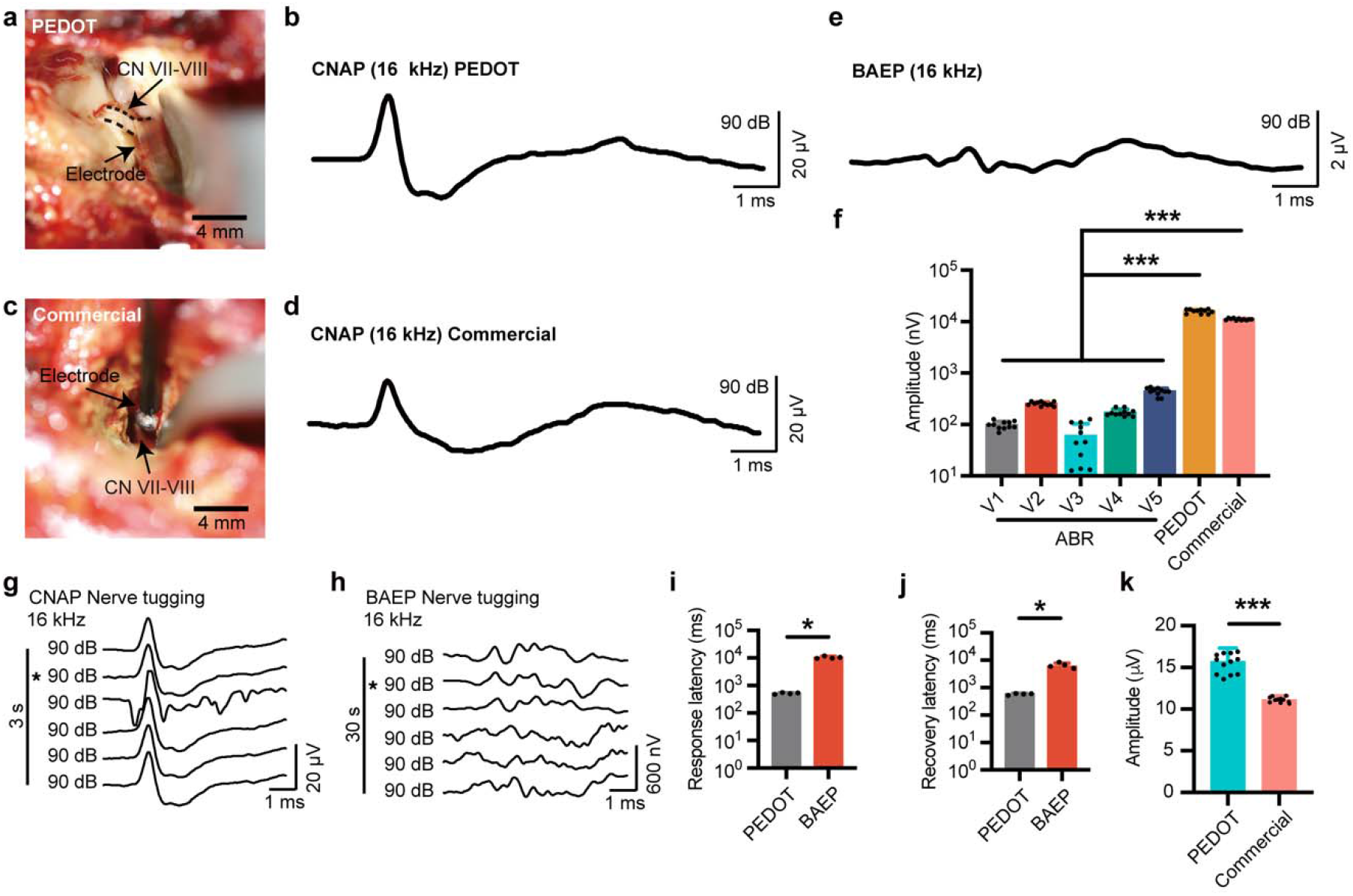
Soft PEDOT electrodes for intra-operative CNAP monitoring. **a,** Photograph of a soft PEDOT electrode wrapping around the facial-acoustic nerve complex for CNAP recording. **b,** CNAP waveforms recorded from the soft PEDOT electrode. **c,** Photograph of a commercial ball-point electrode for CNAP recording. **d,** CNAP waveforms recorded from the conventional metal ball-point electrode. **e,** Control BAEP signals for comparison with the CNAP signals. **f,** Representative auditory monitoring data from BAEP and CNAP. The *P* values for comparison of the amplitudes are as follows: for the BAEP (*n* = 11) versus the PEDOT electrode (*n* = 11), V1 versus the PEDOT, *P* < 0.001; V2 versus the PEDOT, *P* < 0.001; V3 versus the PEDOT, *P* < 0.001; V4 versus the PEDOT, *P* < 0.001; V5 versus the PEDOT, *P* < 0.001. For the BAEP (*n* = 11) versus the commercial electrode (*n* = 11), V1 versus the commercial, *P* < 0.001; V2 versus the commercial, *P* < 0.001; V3 versus the commercial, *P* < 0.001; V4 versus the commercial, *P* < 0.001; V5 versus the commercial, *P* < 0.001. **g-h,** Variation of CNAP (**g**) and BAEP (**h**) waveform during nerve tugging. * denotes the time point of nerve tugging. **i.**Response latencies of BAEP and CNAP after nerve tugging (*n* = 4, *P* = 0.029). **j.** Recovery latencies of BAEP and CNAP after physical nerve tugging (*n* = 4, *P* = 0.029). **k**, Amplitudes of CNAP values recorded using PEDOT and commerical metal electrodes (*n* = 11, *P* < 0.001). All error bars denote s.d. \**P* < 0.05; \*\**P* < 0.01; \*\*\**P* < 0.001; unpaired, two-tailed Student’s t-test was used for **f, i, j,** and **k**. CNAP: Cochlear nerve action potentials; BAEP: Brainstem auditory evoked potentials.

Furthermore, to underscore the advantages of our soft electrode in recording CNAP, we also used a clinically available ball-point probe for the same application. Because of the poor electrode/nerve contacts, the ball-point electrode can only yield ~10 μV for the amplitude (**Fig. 3k**). More importantly, the ball-point electrode was unable to properly maintain stable contacts with the nerve even under mild tugging actions, making it impossible for CINM during real surgical procedures (Fig. S9). To further demonstrate the unique property of PEDOT as the electrode material, we also prepared stretchable micro-cracked gold (Au) electrodes as a control^36^. Although we are able to successfully implant the soft Au electrode around the nerve using the same wrapping method, the CNAP signals recorded using the Au electrode could only reach ~70%of the PEDOT one (Fig. S10). This may likely be due to the higher impedance at the metal/nerve interface^37^. Besides the improved static state signal qualities, the PEDOT electrode also substantially outperformed the Au electrode during dynamic activities. Upon tugging the nerve, while both PEDOT and Au electrodes showed very low response latencies (Fig. S11a-c), and the Au electrode was also observed to require a much longer time to recover (0.58 s vs 1 s, shown in Fig. S11d).

In addition to the long recovery period, the Au electrode also experienced substantial performance drops after the tugging event, i.e., the post-tugging signal amplitude was only ~44.5% of the pre-tugging one (Fig. S11e). This issue is expected to be even more severe in the real-life microsurgery procedure, where multiple unintended tugging motions can occur concurrently. To mimic the possible incidents throughout the surgery, we tugged on the nerve every 30 mins during a 2-hour span and monitored the CNAP signals recorded by both PEDOT and Au electrodes. In this case, the CNAP amplitude recorded using Au showed substantial decays over time and even failed to capture any nerve responses after 2 hours, whereas the PEDOT electrode was able to still consistently record CNAP with nearly constant waveforms (**Fig. 4a-f**). Using optical microscopy to investigate the microstructures of the electrode after the entire surgery, we observed that the Au electrode experienced obvious delamination and wear at the nerve contact sites, which may likely be due to the interfacial micro-motions during the nerve tugging. On the other hand, because of the covalent linkage between PEDOT and the elastomeric substrate (styrene-butadiene-styrene, SBS) during the radical initiated crosslinking reaction, the PEDOT layer remained robustly adhered to the underlying substrate, giving rise to its excellent tolerance to external scratches and frictions (**Fig. 4g-h**).

**Fig. 4.**
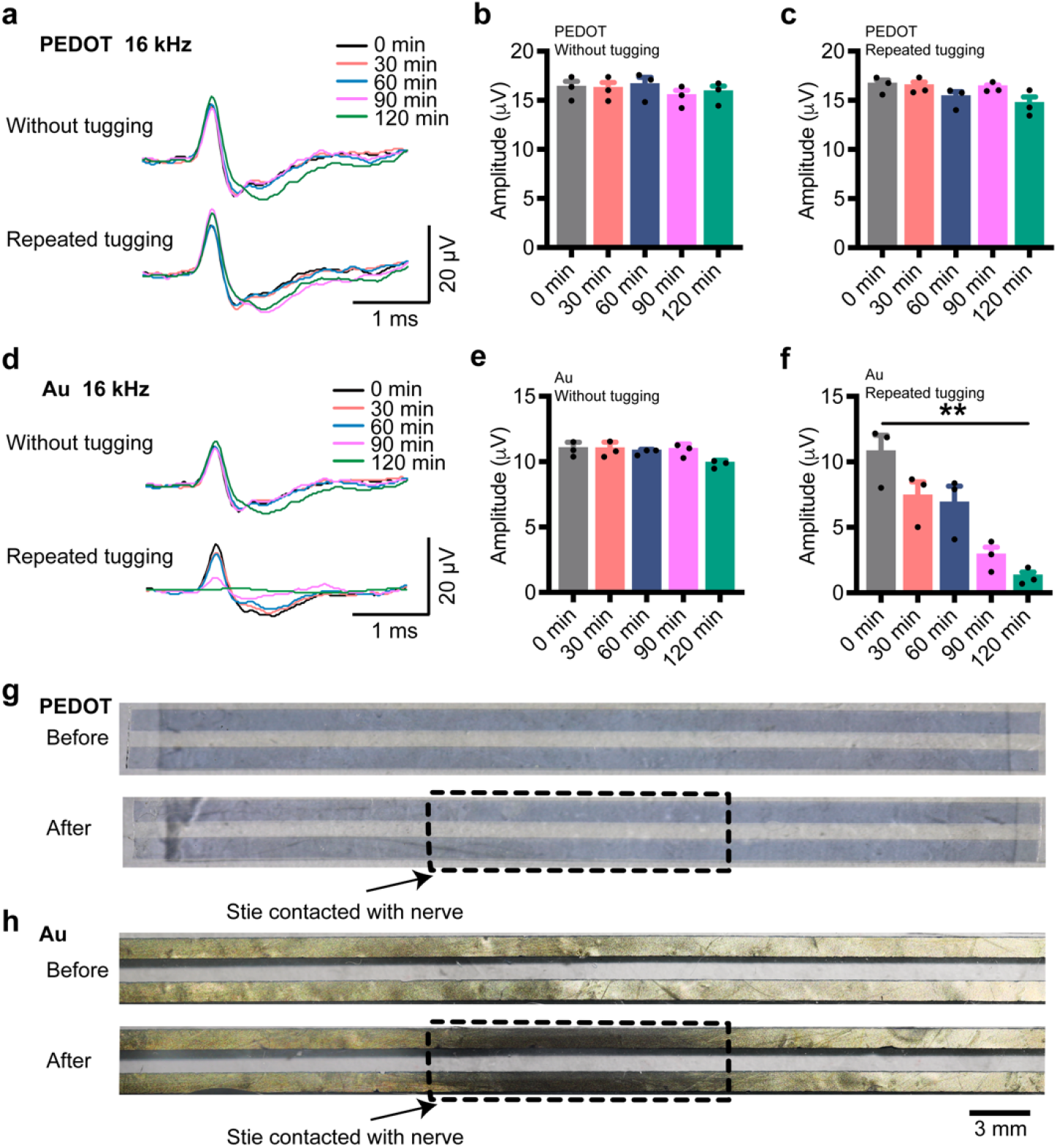
Soft PEDOT electrode for reliable and consistent long-term CNAP monitoring. **a-c,** Continuous intra-operative CNAP monitoring by a soft PEDOT electrode with and without nerve tugging during a 120-min span (*n* = 3). *P* values for comparing CNAP amplitudes without tugging are as follows: for the minute 0 versus minute 30, *P* = 0.700; for the minute 0 versus minute 60, *P* = 0.990; for the minute 0 versus minute 90, *P* = 0.700; for the minute 0 versus minute 120, *P* = 0.700. *P* values for comparing CNAP amplitudes with tugging are as follows: for the minute 0 versus minute 30, *P* = 0.990; for the minute 0 versus minute 60, *P* = 0.400; for the minute 0 versus minute 90, *P* = 0.990; for the minute 0 versus minute 120, *P* = 0.200. **d-f,** Continuous intra-operative CNAP monitoring by an Au electrode with and without nerve tugging during a 120-min span (*n* = 3). *P* values for comparing CNAP amplitudes without tugging are as follows: for the minute 0 versus minute 30, *P* = 0.990; for the minute 0 versus minute 60, *P* = 0.700; for the minute 0 versus minute 90, *P* = 0.990; for the minute 0 versus minute 120, *P* = 0.100. *P* values for comparing CNAP amplitudes with tugging are as follows: for the minute 0 versus minute 30, *P* = 0.400; for the minute 0 versus minute 60, *P* = 0.200; for the minute 0 versus minute 90, *P* = 0.100; for the minute 0 versus minute 120, *P* = 0.002. **g**, Microscope images showing minimal changes of surface apperance of the soft PEDOT electrode after the 120-min CNAP monitoring. **h**, Microscope images showing significant damage of the Au electrode with Au delaminated from the substrate after contacting with the nerve and undergoing micro-motions during the tugging activities during surgery. All error bars denote s.d. \**P* < 0.05; \*\**P* < 0.01; \*\*\**P* < 0.001; unpaired, two-tailed Student’s t-test was used for **b, c, e**, and **f**. CNAP: Cochlear nerve action potentials; CINM: Continuous intra-operative neurophysiological monitoring.

## Hearing preservation without nerve tissue damage by PEDOT

Besides the excellent electrical properties of PEDOT to enable stable recording of CNAP signals, another desired aspect of the electrode for CINM is to have minimal invasiveness for nerve function preservation. Due to the low modulus of SBS elastomer, our soft electrode can be safely integrated with the nerve tissue without causing noticeable hearing loss, as evidenced by the stable BAEP waveforms and minimal changes in decibel threshold for sound detection (**Fig. 5,** Fig. S11a-b). In contrast, when using a stiff polyimide as the substrate, its large modulus mismatch induced considerable changes in BAEP signal amplitude and latency after the tugging motions (**Fig. 5d-g,** Fig. S11c-d). The lowest decibel levels in which the BAEP waveform can be detected also changed from 35 dB to 40 dB (**Fig. 5h**).

**Fig. 5.**
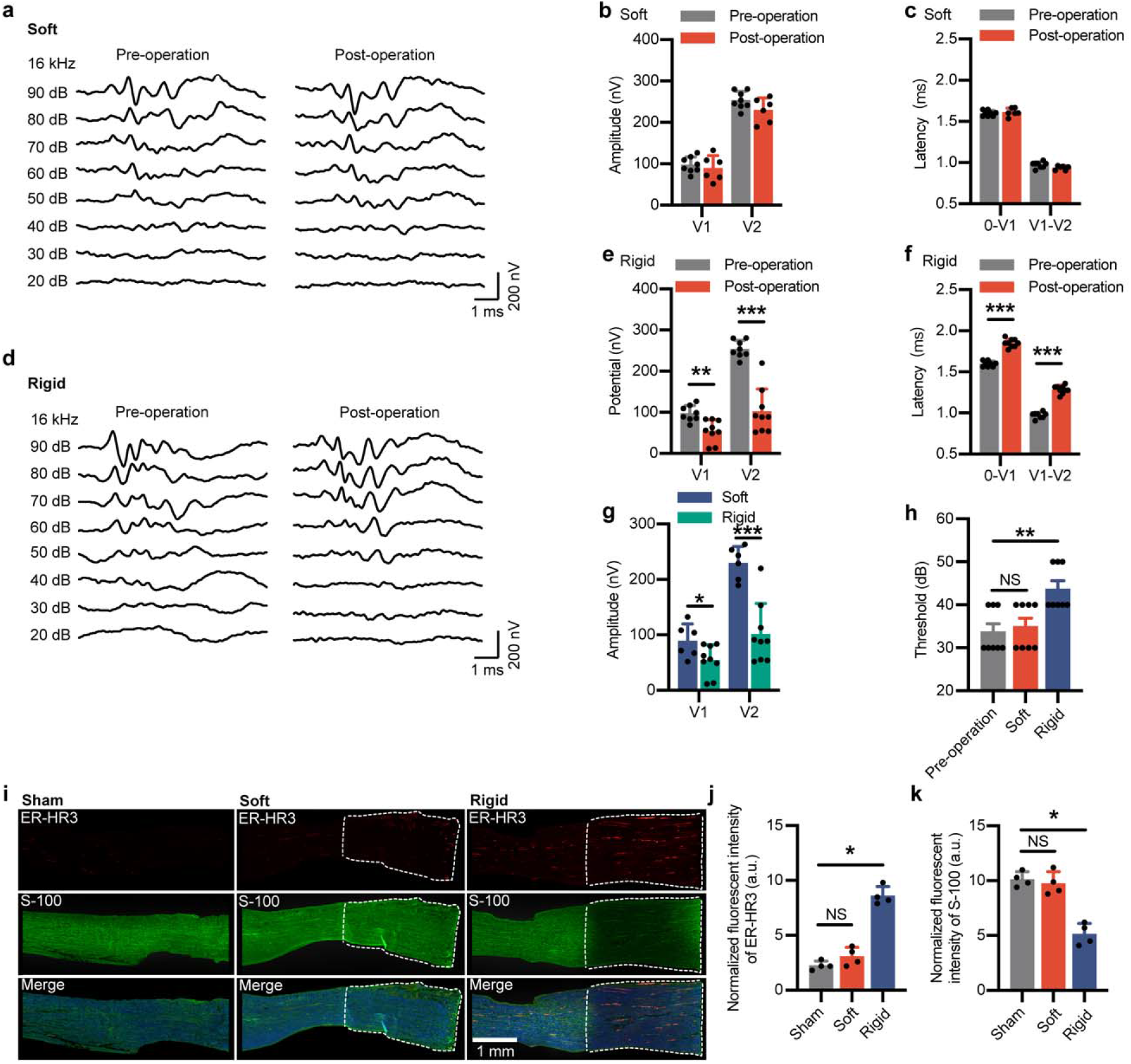
Hearing preservation with minmal nerve tissue damage induced by the soft PEDOT electrode. **a,** BAEP waveforms of the facial-cochlear nerve complex before and after wrapping of a soft PEDOT electrode. **b**, Comparison of BAEP amplitude before and after wrapping of the soft PEDOT electrode. *P* values for comparing the BAEP amplitudes are as follows: V1, pre-operation (*n* = 8) versus post-operation (*n* = 6), *P* = 0.555; V2, pre-operation (*n* = 8) versus post-operation (*n* = 6), *P* = 0.103. **c**, Comparison of BAEP latencies before and after wrapping of the soft PEDOT electrode. *P* values for comparing the BAEP latencies are as follows: 0 - V1, pre-operation (*n* = 8) versus post-operation (*n* = 6), *P* = 0.576; V1 - V2, pre-operation (*n* = 8) versus post-operation (*n* = 6), *P* = 0.070. **d**, BAEP waveforms of the facial-cochlear nerve complex before and after wrapping of a rigid electorde. **e**, Comparison of BAEP amplitudes before and after wrapping of a rigid electorde. *P* values for comparing the BAEP amplitudes are as follows: V1, pre-operation (*n* = 8) versus post-operation (*n* = 9), *P* = 0.002; V2, pre-operation (*n* = 8) versus post-operation (*n* = 9), *P* < 0.001. **f**, Comparison of BAEP latencies before and after wrapping of a rigid electorde. *P* values for comparing the BAEP latencies are as follows: 0 - V1, pre-operation (*n* = 8) versus post-operation (*n* = 9), *P* < 0.001; V1 - V2, pre-operation (*n* = 8) versus post-operation (*n* = 9), *P* < 0.001. **g**, Comparison of BAEP amplitudes after wrapping of a soft PEDOT electrode and a rigid electorde. *P* values for comparing the BAEP amplitudes are as follows: V1, PEDOT electorde (*n* = 8) versus rigid electorde (*n* = 9), *P* = 0.038; V2, PEDOT electorde (*n* = 8) versus rigid electorde (*n* = 9), *P* < 0.001. **h**, Hearing threshold pre-operation, after wrapping of soft PEDOT and rigid electrodes. *P* values for comparing the hearing thresholds are as follows: pre-operation (*n* = 8) versus PEDOT electrode (*n* = 8), *P* = 0.990; pre-operation (*n* = 8) versus rigid electrode (*n* = 8), *P* = 0.009. **i**, Longitudinal-section slice of the facial-cochlear nerve complex labeled by an inflammatory biomarker (macrophage/monocyte) and a Schwann cell biomarker (S-100) for soft PEDOT electrode, rigid electrode and sham control. **j-k**, Comparison of normalized fluorescence intensities from ER-HR3 (**j**) and S-100 (**k**) for soft PEDOT electrode, rigid electrode and sham control. *P* values for comparing ER-HR3 intensities are as follows: sham (*n* = 4) and PEDOT electrode (*n* = 4), *P* = 0.200; sham (*n* = 4) and rigid electrode (*n* = 4), *P* = 0.029. *P* values for comparing the S-100 intensities are as follows: sham (*n* = 4) and PEDOT electrode (*n* = 4), *P* = 0.686; sham (*n* = 4) and rigid electrode (*n* = 4), *P* = 0.029. All error bars denote s.d. \**P* < 0.05; \*\**P* < 0.01; \*\*\**P* < 0.001; NS, not significant; unpaired, two-tailed Student’s t-test was used for **b, c, e, f, g, h, j**, and **k**. BAEP: Brainstem auditory evoked potentials.

Furthermore, as compared to the sham control, confocal microscopy images of immunostained nerve tissue slices revealed no significant differences in fluorescence intensities of macrophage/monocyte markers (ER-HR3) in the nerve after tugging the soft electrode. On the contrary, the implanted rigid electrode caused significant inflammatory responses (**Fig. 5i-j**). Additionally, we investigated the nerve injury by quantifying S-100, a hallmark of Schwann cells. When comparing in between the nerve with the soft electrode and the sham control, there was no statistically significant differences in S-100 expression levels. However, the rigid electrode was observed to cause substantial damage to the nerve with a clear boundary around the injury site with reduced S-100 expression (**Fig. 5i, 5k**). To verify the long-term biocompatibility of our device, we implanted both soft and rigid electrodes onto sciatic nerves in freely moving mice for two weeks (Fig. S12a). Similar to results obtained from the short-term tugging experiment, the rigid electrode again induced substantially elevated immune responses, whereas the soft device remained inactive (Fig. S12b).

## Optimal prognosis was achieved with PEDOT monitoring

To evaluate the impact of CINM enabled by the soft and stretchable PEDOT electrode, we monitored the prognosis in tumor-bearing rats after microsurgery (Fig. S14). To reproduce the schwannoma animal model, we first implanted tumor cells into the rat sciatic nerve^38,39^, and then investigated EMG and gait changes before and after tumor resection. During the microsurgical operation, since the tumor is fully integrated with the nerve itself, it is impossible to distinguish them under the naked eye. Even with intermittent monitoring of neurophysiological signals using the commercial ball-point electrode, the micro-scissors still inevitably caused sharp nerve transection during the tumor excision. As a result, EMG signals collected from the same rat showed reduced amplitude at 4 weeks after surgery (**Fig. 6a-c,** Fig. S16). In contrast, with the PEDOT electrode to perform CINM, we were able to capture evoked neural responses immediately upon mechanical contact, therefore preventing further damages by the surgical manipulation. In this instance, because of the undamaged nerve structure following the tumor removal, the EMG amplitude was significantly higher after the surgery (**Fig. 6d-f,** Fig. S16). We also evaluated the gait patterns of tumor bearing rats using kinematic analysis (**Fig. 6g,** Video S3). Using only commercial electrodes for intermittent monitoring, the rat was observed to experience worse paw function with whole-foot placement on the ground and dragging of their toes (**Fig. 6h,** Fig. S17b, Video S4), a typical symptom of serious sciatic nerve dysfunction. In contrast, as aided by CINM using PEDOT, the rat showed significantly better gait quality after the surgery^40^, as evidenced by the improved paw placement (**Fig. 6i,** Fig. S17a, Supplementary Video 4).

**Fig. 6.**
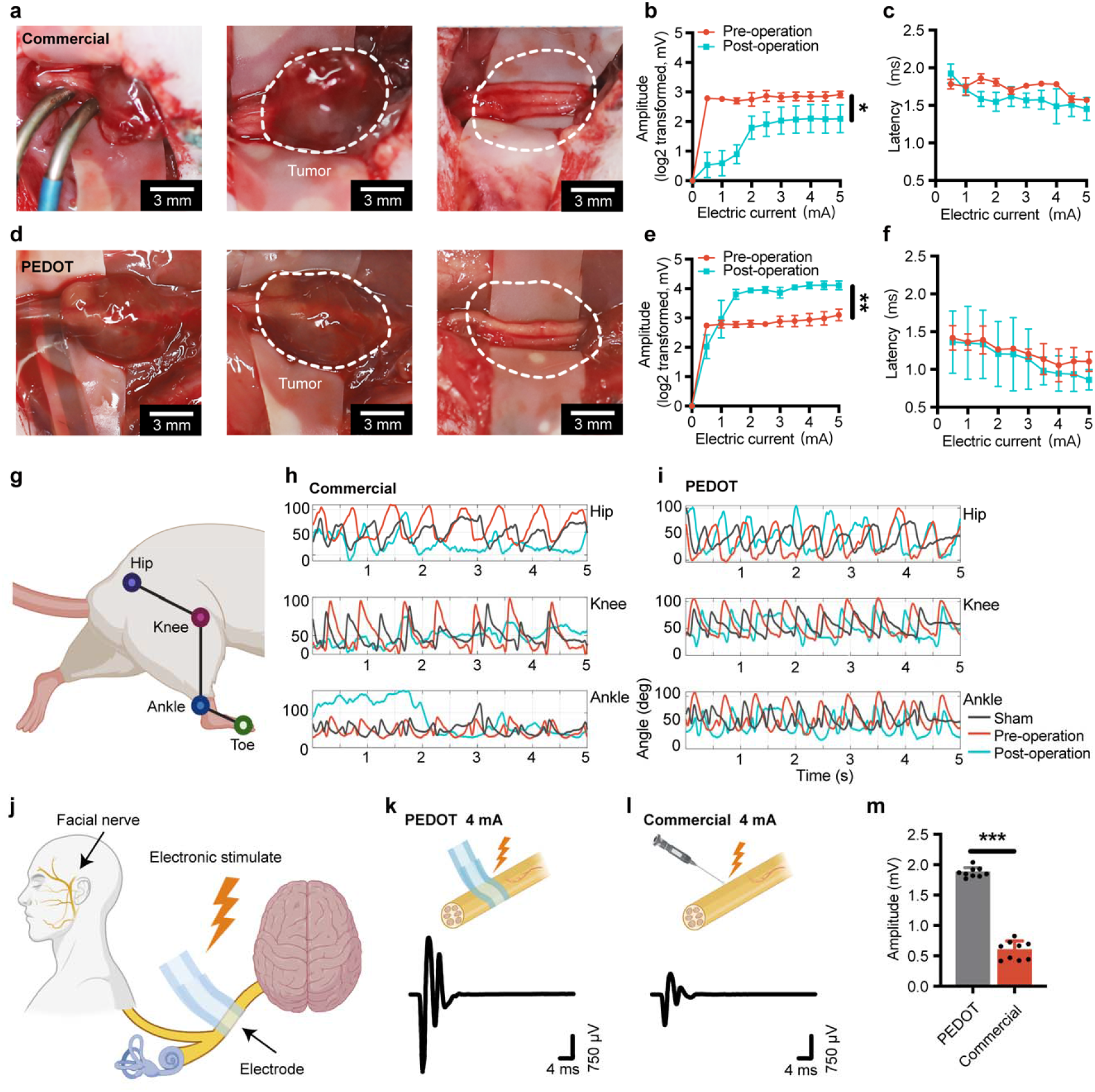
Improved post-operative prognosis using soft PEDOT electrodes. **a,** Photographs showing a commercial metal electrode for intra-operative monitoring, and a sciatic nerve with tumor before after resection. **b-c**, Comparison of evoked electromyographic amplitudes (**b**) and latencies (**c**) before tumor removal and 4 weeks after resection using a commercial metal electrode for intra-operative monitoring (**b**, *n* = 4, *P* = 0.024; **c**, *n* = 4, *P* = 0.354). **d**, Photographs showing a soft PEDOT electrode for intra-operative monitoring, and a sciatic nerve with tumor before after resection. **e-f**, Comparison of evoked electromyographic amplitudes (**e**) and latencies (**f**) before tumor removal and 4 weeks after resection using a commercial metal electrode for intra-operative monitoring (**e**, *n* = 4, *P* = 0.003; **f**, *n* = 4, *P* = 0.387). **g**, Schematic diagram illustrating the hindlimb kinematics during walking. **h-i**, Representative angular movements for the hip, knee and ankle joints for the mouse before and 4 weeks after tumor removal on the sciatic nerve by commercial metal electrode (**h**) and soft PEDOT (**i**) monitoring. **j**, Schematic of neural stimulation for facial nerve evaluation. **k-l**, Schematic and evoked electromyography waveforms by stimulating the facial nerves using a PEDOT electrode (k) and a commercial metal electrode (**l**) at 4 mA. **m**, Comparison of evoked electromyography amplitudes by PEDOT electrode and conventional metal electrode at 4 mA (*n* = 9, *P* < 0.001). All error bars denote s.d. \**P* < 0.05; \*\**P* < 0.01; \*\*\**P* < 0.001; unpaired, two-tailed Student’s t-test was used for **d**. One-way ANOVA with Holm-Sidak’s multiple comparisons test was used for **f, g, i,** and **j**.

Finally, beside recording neurophysiological signals, the low-impedance PEDOT electrode also allows functional stimulation to evaluate neural functions during surgery. This modality is particularly useful to preserve motor nerve functions and preventing post-operative paralysis. For VS surgery, facial nerve paralysis is one of the most clinically significant complications owing to the close proximity between cochlear and facial nerves^41^. In ~44% of individuals with VS, surgical treatment resulted in persistent facial weakness^41,42^. Localized stimulation of the facial nerve is a promising strategy for minimizing the risk of facial nerve injury, as it ensures that downstream muscle actions are undisturbed during the surgical operation (**Fig. 6j**). Using a rabbit model, we first implanted the soft PEDOT electrode to form an intimate contact with the facial-cochlear nerve complex (Video S5). Because of the low interfacial impedance and low modulus of our PEDOT electrode, we could induce noticeable facial muscle movements at an ultralow threshold of 1 mA, without the need to remove or penetrate the epineurium layer. In comparison, the commercial ball-point electrode would require at least 4 mA to elicit noticeable facial movements (Video S6). The amplitude of the EMG signal elicited by the PEDOT electrode was also substantially higher than that induced by the ball-point electrode (**Fig. 6k-m**). In the clinical setting, this highly efficient stimulation functionality will allow the use of the PEDOT electrode to forecast the severity of facial paralysis in patients undergoing VS surgery.

## Conclusion

We have demonstrated that CINM, in combination with advanced micro-neurosurgical techniques, is extremely valuable to prevent neurological injury during surgery. In this work, by developing a clinically deployable bioelectronic device based on soft and low impedance conducting polymers, we successfully achieved *for the first time* continuous recording of near-field action potential with high signal-to-noise ratio and minimal invasiveness during tumor resection surgeries, leading to dramatically reduced post-operative morbidity in pre-clinical animal models. Utilizing this unprecedented CINM modality in conjunction with localized neurostimulation, we further demonstrated our approach to successfully restore normal functions after a variety of neurosurgeries. Moving forward, further development of chronically stable, high-resolution, soft and stretchable electrode arrays with efficient neurostimulation and recording modalities should allow successful applications in more complicated surgical disciplines, and even for long-term and closed-loop disease management.

## Acknowledgments

This work was supported by the National Natural Science Foundation of China (No. 82071996). Part of this work was performed at the Stanford Nano Shared Facilities (SNSF), supported by the National Science Foundation under award ECCS-2026822. We thank Guijun Jia and Xue Zhan for administrative support. We thank Lirui Yang for instrument support of CINM measurements. We also thank Dr. Chuanbao Zhang, Dr. Xi Wang, Dr. Yangyang Wang for their guidance with the project.

## Author Contributions

W.Z., Y.J., D.L., Z.B. and W.J. designed the study. W.Z., Y.J., Q.X., L.C., M.L. performed circuit design and testing. Y.J., J.L., D.Z., Y.W. performed material synthesis and characterizations. W.Z., Q.X., H.Q., Y.Z., W.L., X.W., J.L., X.G., S.M., P.K., L.Y., J.B.T. performed the animal, cell culture experiments and histological staining. W.Z., Y.J., D.L. and Z.B. wrote the manuscript with input from all co-authors.

## Methods

A complete, detailed description of methods can be found in the Supplementary Information.

## References

1 Buckner, J. C. et al. Central nervous system tumors. Mayo Clin Proc 82, 1271–1286, doi:10.4065/82.10.1271 (2007).

2 Horbinski, C., Berger, T., Packer, R. J. & Wen, P. Y. Clinical implications of the 2021 edition of the WHO classification of central nervous system tumours. Nat Rev Neurol, doi:10.1038/s41582-022-00679-w (2022).

3 Miller, K. D. et al. Brain and other central nervous system tumor statistics, 2021. CA Cancer J Clin 71, 381–406, doi:10.3322/caac.21693 (2021).

4 Carlson, M. L. & Link, M. J. Vestibular Schwannomas. N Engl J Med 384, 1335–1348, doi:10.1056/NEJMra2020394 (2021).

5 Goldbrunner, R. et al. EANO guideline on the diagnosis and treatment of vestibular schwannoma. Neuro Oncol 22, 31–45, doi:10.1093/neuonc/noz153 (2020).

6 Sanai, N. & Berger, M. S. Surgical oncology for gliomas: the state of the art. Nat Rev Clin Oncol 15, 112–125, doi:10.1038/nrclinonc.2017.171 (2018).

7 Lapointe, S., Perry, A. & Butowski, N. A. Primary brain tumours in adults. The Lancet 392, 432–446, doi:10.1016/s0140-6736(18)30990-5 (2018).

8 Betka, J. et al. Complications of microsurgery of vestibular schwannoma. Biomed Res Int 2014, 315952, doi:10.1155/2014/315952 (2014).

9 Hirbe, A. C. & Gutmann, D. H. Neurofibromatosis type 1: a multidisciplinary approach to care. Lancet Neurol 13, 834–843, doi:10.1016/S1474-4422(14)70063-8 (2014).

10 Gonzalez, A. A., Jeyanandarajan, D., Hansen, C., Zada, G. & Hsieh, P. C. Intraoperative neurophysiological monitoring during spine surgery: a review. Neurosurg Focus 27, E6, doi:10.3171/2009.8.FOCUS09150 (2009).

11 Rho, Y. J., Rhim, S. C. & Kang, J. K. Is intraoperative neurophysiological monitoring valuable predicting postoperative neurological recovery? Spinal Cord 54, 1121–1126, doi:10.1038/sc.2016.65 (2016).

12 Liu, Y. et al. Intraoperative monitoring of neuromuscular function with soft, skin-mounted wireless devices. NPJ Digit Med 1, doi:10.1038/s41746-018-0023-7 (2018).

13 Langguth, B., Kreuzer, P. M., Kleinjung, T. & De Ridder, D. Tinnitus: causes and clinical management. Lancet Neurol 12, 920–930, doi:10.1016/S1474-4422(13)70160-1 (2013).

14 Watanabe, N. et al. Intraoperative cochlear nerve mapping with the mobile cochlear nerve compound action potential tracer in vestibular schwannoma surgery. J Neurosurg, 1–8, doi:10.3171/2017.12.JNS171545 (2018).

15 Nakatomi, H. et al. Improved preservation of function during acoustic neuroma surgery. J Neurosurg 122, 24–33, doi:10.3171/2014.8.JNS132525 (2015).

16 Piccirillo, E. et al. Intraoperative cochlear nerve monitoring in vestibular schwannoma surgery--does it really affect hearing outcome? Audiol Neurootol 13, 58–64, doi:10.1159/000108623 (2008).

17 Legatt, A. D. Electrophysiology of Cranial Nerve Testing: Auditory Nerve. J Clin Neurophysiol 35, 25–38, doi:10.1097/WNP.0000000000000421 (2018).

18 Yamakami, I., Yoshinori, H., Saeki, N., Wada, M. & Oka, N. Hearing preservation and intraoperative auditory brainstem response and cochlear nerve compound action potential monitoring in the removal of small acoustic neurinoma via the retrosigmoid approach. J Neurol Neurosurg Psychiatry 80, 218–227, doi:10.1136/jnnp.2008.156919 (2009).

19 Yamakami, I., Oka, N. & Yamaura, A. Intraoperative monitoring of cochlear nerve compound action potential in cerebellopontine angle tumour removal. J Clin Neurosci 10, 567–570, doi:10.1016/s0967-5868(03)00143-7 (2003).

20 O’Doherty, J. E. et al. Active tactile exploration using a brain–machine–brain interface. Nature 479, 228–231, doi:10.1038/nature10489 (2011).

21 Betzel, R. F. et al. Structural, geometric and genetic factors predict interregional brain connectivity patterns probed by electrocorticography. Nature Biomedical Engineering 3, 902–916, doi:10.1038/s41551-019-0404-5 (2019).

22 Miyazaki, H. & Caye-Thomasen, P. Intraoperative Auditory System Monitoring. Adv Otorhinolaryngol 81, 123–132, doi:10.1159/000485577 (2018).

23 Yuk, H., Lu, B. & Zhao, X. Hydrogel bioelectronics. Chem Soc Rev 48, 1642–1667, doi:10.1039/c8cs00595h (2019).

24 Paulsen, B. D., Tybrandt, K., Stavrinidou, E. & Rivnay, J. Organic mixed ionic-electronic conductors. Nat Mater 19, 13–26, doi:10.1038/s41563-019-0435-z (2020).

25 Helbing, D. L., Schulz, A. & Morrison, H. Pathomechanisms in schwannoma development and progression. Oncogene 39, 5421–5429, doi:10.1038/s41388-020-1374-5 (2020).

26 Ammoun, S. & Hanemann, C. O. Emerging therapeutic targets in schwannomas and other merlin-deficient tumors. Nat Rev Neurol 7, 392–399, doi:10.1038/nrneurol.2011.82 (2011).

27 Matthies, C. & Samii, M. Management of 1000 vestibular schwannomas (acoustic neuromas): clinical presentation. Neurosurgery 40, 1–9; discussion 9-10, doi:10.1097/00006123-199701000-00001 (1997).

28 Kirchmann, M. et al. Ten-Year Follow-up on Tumor Growth and Hearing in Patients Observed With an Intracanalicular Vestibular Schwannoma. Neurosurgery 80, 49–56, doi:10.1227/NEU.0000000000001414 (2017).

29 Propp, J. M., McCarthy, B. J., Davis, F. G. & Preston-Martin, S. Descriptive epidemiology of vestibular schwannomas. Neuro Oncol 8, 1–11, doi:10.1215/S1522851704001097 (2006).

30 Ochal-Choinska, A., Lachowska, M., Kurczak, K. & Niemczyk, K. Audiologic prognostic factors for hearing preservation following vestibular schwannoma surgery. Adv Clin Exp Med 28, 747–757, doi:10.17219/acem/90768 (2019).

31 Zhou, W. et al. A Novel Imaging Grading Biomarker for Predicting Hearing Loss in Acoustic Neuromas. Clin Neuroradiol, doi:10.1007/s00062-020-00938-7 (2020).

32 Someya, T., Bao, Z. & Malliaras, G. G. The rise of plastic bioelectronics. Nature 540, 379–385, doi:10.1038/nature21004 (2016).

33 Khodagholy, D. et al. NeuroGrid: recording action potentials from the surface of the brain. Nat Neurosci 18, 310–315, doi:10.1038/nn.3905 (2015).

34 Jiang, Y. et al. Topological supramolecular network enabled high–conductivity, stretchable organic bioelectronics. Science 375, 1411–1417, doi:10.1126/science.abj7564 (2022).

35 Yamakami, I., Ito, S. & Higuchi, Y. Retrosigmoid removal of small acoustic neuroma: curative tumor removal with preservation of function. J Neurosurg 121, 554–563, doi:10.3171/2014.6.JNS132471 (2014).

36 Lacour, S. P., Chan, D., Wagner, S., Li, T. & Suo, Z. Mechanisms of reversible stretchability of thin metal films on elastomeric substrates. Applied Physics Letters 88, 204103, doi:10.1063/1.2201874 (2006).

37 Liu, Y. et al. Soft and elastic hydrogel-based microelectronics for localized low-voltage neuromodulation. Nat Biomed Eng 3, 58–68, doi:10.1038/s41551-018-0335-6 (2019).

38 Gao, X. et al. Anti-VEGF treatment improves neurological function and augments radiation response in NF2 schwannoma model. Proc Natl Acad Sci U S A 112, 14676–14681, doi:10.1073/pnas.1512570112 (2015).

39 Wu, L. et al. Losartan prevents tumor-induced hearing loss and augments radiation efficacy in NF2 schwannoma rodent models. Sci Transl Med 13, 4816, doi:10.1126/scitranslmed.abd4816 (2021).

40 de Medinaceli, L., Freed, W. J. & Wyatt, R. J. An index of the functional condition of rat sciatic nerve based on measurements made from walking tracks. Experimental Neurology 77, 634–643, doi:10.1016/0014-4886(82)90234-5 (1982).

41 Leong, S. C. & Lesser, T. H. A national survey of facial paralysis on the quality of life of patients with acoustic neuroma. Otology & neurotology 36, 503–509 (2015).

42 Owusu, J. A., Stewart, C. M. & Boahene, K. Facial Nerve Paralysis. Med Clin North Am 102, 1135–1143, doi:10.1016/j.mcna.2018.06.011 (2018).

