## Supplemental Figures for "Soft and stretchable organic bioelectronics for continuous intra-operative neurophysiological monitoring during microsurgery"

### Methods

**Fabrication of PEDOT:PSS electrode.** Poly(3,4-ethylenedioxythiophene) polystyrene sulfonate (PEDOT:PSS, Orgacon™ ICP 1050) was provided by Agfa-Gevaert N.V as a surfactant-free aqueous dispersion with 1.1 wt% solid content. Polyrotaxane (PR) based supramolecular crosslinker was synthesized based on a previous report. Styrene-butadiene-styrene (SBS) D1102 was provided by Kraton as the elastomeric substrate.

For the SBS substrate preparation, clean glass substrates were first treated with $O_2$ plasma at 150 W for 30 s using a Technics Micro-RIE Series 800. Next, a dextran (from *Leuconostoc spp.*, Mr ~100,000 Sigma-Aldrich) aqueous solution (5 wt%) was spin coated at 500 rpm for 15 s as the sacrificial layer. The SBS substrate was then prepared by drop casting 80 mg $mL^{-1}$ D1102, 2.4 mg $mL^{-1}$ pentaerythritol tetrakis(3-mercaptopropionate) (PETMP, Sigma-Aldrich), and 0.08 mg mL phenylbis(2,4,6-trimethylbenzoyl)phosphine oxide (BAPO, Sigma-Aldrich) in toluene (Fisher) and subsequent drying at room temperature overnight. The as-dried substrate was then crosslinked under ultraviolet (UV) illumination for 10 min using an ELC-500 UV curing chamber (Electro-Lite, 30 mW $cm^{-2}$) in glovebox and baked at 150 ºC for 10 min under the nitrogen atmosphere. The final substrate thickness was ~100 μm to ensure easy handling and low bending stiffness.

For the PEDOT electrode layer, 55 mg of PR together with 1 mg of photo initiator, lithium phenyl-2,4,6-trimethylbenzoylphosphinate (LAP, Sigma-Aldrich), and 5 μL of fluorosurfactant (Zonyl FS-30, Sigma-Aldrich) were added to 1 mL of PEDOT:PSS aqueous dispersion (Orgacon™ ICP 1050, 1.1 wt%) and mixed using a vortex mixer. After filtering using nylon syringe filters (pore size: 1 μm, Whatman), the mixture would be spin coated onto the crosslinked SBS substrate at 500 rpm for 15 s. The spin coated film would be later photo-patterned by UV exposure for 2 min using a Spectrum 100 UV curing system (American Ultraviolet, 25 mW $cm^{-2}$) before rinsing in water for development. The as-patterned PEDOT electrode was further dipped in methanol (Fisher Scientific) for 1 min to improve the conductivity before use.

**Cell lines.** Rat RT4-D6P2T Schwann cells were maintained in 10% Schwann cell medium containing Schwann cell growth supplement (SCGS, ScienCell, Carlsbad, CA)[1,2].

**Animal experiments.** All animal procedures were performed following the guidelines of Public Health Service Policy on Humane Care of Laboratory Animals and approved by the Institutional Animal Care and Use Committee of the Capital Medical University, Beijing Tiantan Hospital.

**Intra-operative auditory monitoring.** To replicate the retrosigmoid craniotomy on human VS patients, we performed the whole process of retrosigmoid craniotomy in a rabbit model. the rabbit was anesthetized with 2% isoflurane in balanced oxygen.

Brainstem auditory evoked potentials (BAEP) monitoring. Under sterile conditions with external body warming, the speaker is placed 10 cm from the operated side ear of the rabbit. For hearing monitoring, 3 subdermal electrodes were used: ground, reference, and active electrode, which were placed 2-3 mm under the skin of the rabbit head. Insert the active electrode subdermally into the forehead. Insert the reference electrode below the pinna of the right ear, and the ground electrode below the contralateral (left) ear. Sounds are presented and BAEPs are recorded in a free field condition with pure tone (BioSig III, TDT, USA) and hardware (TDT RZ6 and TD speakers, USA) are used for BAEP recording[3].

Cochlear nerve action potentials (CNAP) monitoring. Under sterile conditions with external body warming, a linear vertically oriented occipital incision and a craniotomy just posterior and inferior to the sigmoid and transverse sinuses, respectively, were used in the surgery of rabbit. The dura was then opened, exposing the facial-acoustic nerve complex with the cerebellum retracted. The facial-acoustic nerve complex was either (1) inserted into the commercial metal electrode or (2) wrapped with the PEDOT electrode for auditory monitoring[4]. At the end of surgery, rabbits were killed with carbon dioxide gas followed by decapitation for proper disposal. The facial-acoustic nerve complexes were collected for analysis.

***In vivo* facial nerve electrical stimulation.** For facial nerve stimulation, the rabbit was anesthetized with 2% isoflurane in balanced oxygen. Under sterile conditions with external body warming, the facial-acoustic nerve complex was wrapped with the PEDOT electrode or implanted with commercial metal electrode for facial nerve electrical stimulation. For the whisker movement experiment, 20 Hz, biphasic charge balanced rectangular voltage pulses (200 μs pulse width) were applied by using a function generator (Neuron-spectrum-4/EPM, Neurosoft, Russia). The whisker response was recorded using a digital camera. EMG was used to record muscle activity during electrical stimulation of the facial nerve. Two needle electrodes were used to penetrate the muscles, and the electrodes were connected to a signal acquisition system (Neuro-MEPw, Neurosoft, Russia). Various electric currents (from 0.5 mA to 5 mA) were used to stimulate the facial nerve while EMG traces was recorded. The rabbits were killed with carbon dioxide gas immediately after the experiment.

**Implantation of PEDOT electrode on rat sciatic nerves.** Wistar rats was anesthetized with 2% isoflurane in balanced oxygen. Under sterile conditions with external body warming, a 3-cm incision was made on the right thigh. About 1 cm of the sciatic nerve proximal to the tibial and peroneal bifurcation was either (1) inserted into the commercial metal electrode or (2) wrapped with the PEDOT electrode for CINM[4]. 2 weeks after surgery, rats were killed with carbon dioxide gas followed by decapitation for proper disposal. Sciatic nerves were collected for further analysis.

**Sciatic nerve tumor model.** The soft peripheral nervous system is a desirable organ system for soft and conductive polymer-based electronic devices[5,6]. To reproduce the schwannoma animal model, we implanted tumor cells into the rat sciatic nerve[7,8]. RT4-D6P2T schwannoma cells were implanted in 8-12 weeks old immune competent Wistar rats. A total of 10 μl of tumor cell suspension ($1x10^6$ cells) was injected slowly (over 45-60 seconds) under the sciatic nerve sheath using Hamilton syringe to prevent leakage. 12 days after implantation, tumor size was measured in diameter and EMG was monitored to ensure that there were no differences between rats in different experimental groups. The tumors were resected under microscope (M525-F20, Leica Microsystems, Germany). Samples were collected for further staining.

**Electrical stimulation of the sciatic nerve.** The rat was anesthetized with 2% isoflurane in balanced oxygen. Under sterile conditions with external body warming, the sciatic nerve was wrapped with the PEDOT electrode or implanted with commercial metal electrode for sciatic nerve electrical stimulation. For the leg movement experiment, 20 Hz, biphasic charge balanced rectangular voltage pulses (200 μs pulse width) were applied by using a function generator (Neuron-spectrum-4/EPM, Neurosoft, Russia). EMG was used to record muscle activity during electrical stimulation of the sciatic nerve. Two needle electrodes were used to penetrate the muscles, and the electrodes were connected to a signal acquisition system (Neuro-MEPw, Neurosoft, Russia). Various electric currents (from 0.5 mA to 5 mA) were used to stimulate the sciatic nerve while EMG traces was recorded before tumor resection and 4 weeks after tumor removal. The rats were killed immediately after the experiment.

**Behavior analysis.** Animals were divided into matched groups and behavioral testing was performed by double-blinded individuals. Animals were trained on 3 separate days before recording baseline behavior. After tumor implantation, gait analysis was assessed weekly. A stick diagram decomposition of hindlimb movements was plotted based on the video, in which the rats are walking on a treadmill. Mice were recorded in the sagittal plane while walking on a treadmill[9]. A high-speed camera (240 frames/s; Fastec IL3-100, San Diego,

CA, USA) was used to record the animals from the side while they walked steadily for 10–12 step cycles. Rats were recorded at a speed of 20 cm/s (cps). Videos of the mice walking on the treadmill were analyzed by ImageJ and R. In brief, the intensity threshold was adjusted for each video so that only the reflective markers were visible. This resulted in a video for which only the five markers on the animal's leg were visible in each frame. The X and Y coordinates, in pixels, of these markers were then tracked frame by frame for the duration of the recording, allowing stick model reconstructions of the leg to be made[10].

**Immunohistology and immunostaining.** Biocompatibility studies were performed to compare nerve bundle damage and immune responses at the implantation site for the soft PEDOT electrode, the rigid electrode, and the sham control. In the short-term biocompatibility study, four rabbits were used for each group. After monitoring, the rabbits were killed. In the long-term biocompatibility study, four rats were used for each group. The rats were killed 2 weeks after implantation. The nerves with implants were fixed in a PBS solution with 4 wt% formaldehyde for 24 h. Fixed nerves were first transferred to a sucrose solution (30%) overnight, then transferred to a sucrose–Cryo-OCT compound (VWR International) solution for 8 h and finally transferred to Cryo-OCT compound and frozen at −80 °C. The frozen samples were sectioned into 10-μm tissue slices using a Leica CM1950 cryosectioning instrument (Leica Microsystems, Germany).

ER-HR3 (Macrophage/monocytes) and S-100 in the tissue slices were stained for imaging. The slices were washed with PBS three times and then permeabilized by 0.1% Triton-X 100 in PBS for 15 min. After washing with PBS solution, the samples were incubated in blocking solution (3% bovine serum albumin, 0.1% Triton-X 100 in PBS) for 45 min. The samples were then co-stained with 1:100 anti-ER-HR3 (ab59697, Abcam) and 1:50 anti-S-100 (ab218515, Abcam) in blocking solution overnight at 4°C. The samples were washed with PBS and stained with secondary antibody anti-mouse IgG H&L (Alexa Fluor 488, ab150113) and anti-rat IgG H&L (Alexa Fluor 647, ab150167). Confocal images of the samples were taken using an Inverted Zeiss LSM 780 multiphoton laser scanning microscope. The fluorescence intensity and inflammatory tissue area were normalized to the mean value of the sham control. Unpaired, two-tailed t-tests were performed using Prism.

**Statistical analysis.** R, Prism and Excel were used for all statistical analyses. All replicate numbers, error bars, *P* values and statistical tests are indicated in figure legends.

**Reporting Summary.** Further information on experimental design is available in the Nature Research Reporting Summary linked to this article.

**Data availability.** The authors declare that all data supporting the findings of this study are available within the paper and its Supplementary Information.

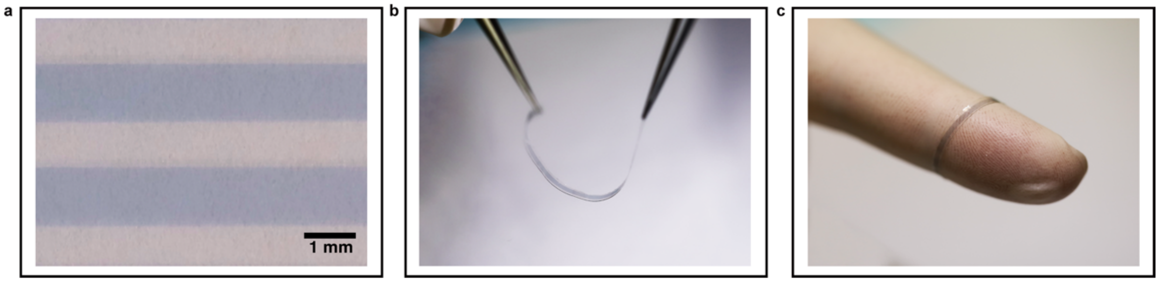


**Supplementary Fig. 1 | Photographs of soft PEDOT electrode. a,** Microscope image of the patterned stretchable PEDOT electrode based on the polyrotaxane additive. **b-c,** Soft and stretchable PEDOT electrode can be easily bent (**b**) and wrap around tissues (**c**).

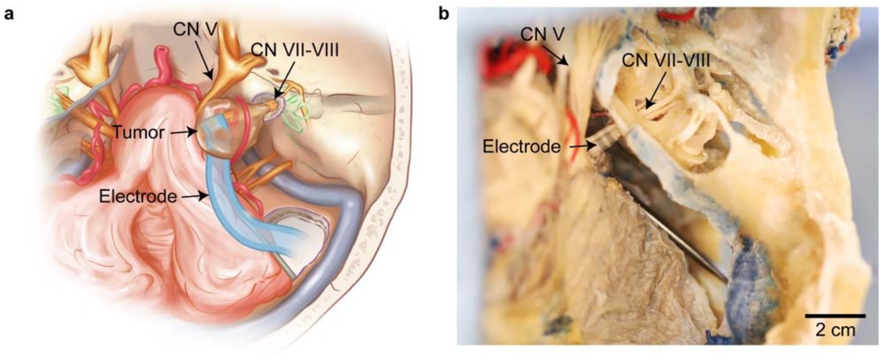


**Supplementary Fig. 2 | Schematic (a) and photograph (b) of the PEDOT electrode wrapped around the facial-acoustic nerve complex.** The PEDOT devices was implanted in a human cadaver skull using the retrosigmoid approach from superior view of axial cross section.

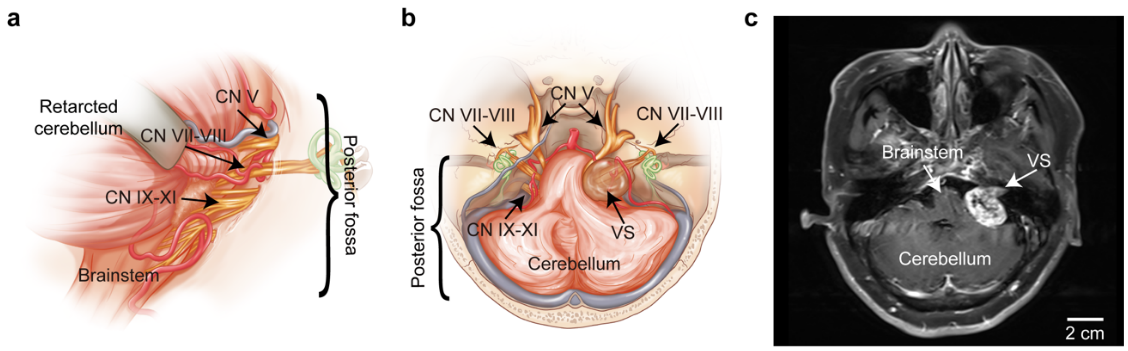


**Supplementary Fig. 3 | Microanatomy and structures affected by VS. a,** Schematic showing the relevant microanatomy of the posterior fossa. **b-c,** Schematic (**b**) and MRI image (**c**) showing the effect of tumor growth on the adjacent cranial nerves, the brainstem, and the cerebellum. VS characteristically arise within the internal auditory canal, from one of the two vestibular divisions of the vestibulocochlear nerve. VS: Vestibular schwannomas.

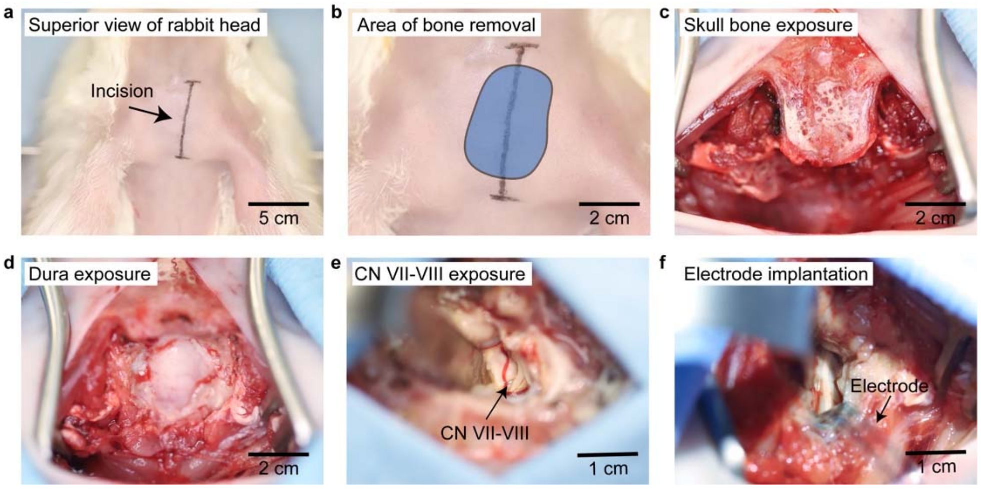


**Supplementary Fig. 4 | The whole process of retrosigmoid craniotomy in a rabbit model. a,** A linear, vertically oriented occipital incision was used in the surgery. **b-c,** Bones that are just posterior and inferior to the sigmoid and transverse sinuses were removed, respectively. **d,** After the bone was removed, the dura was exposed. **e,** The dura was then opened, exposing the facial-acoustic nerve complex with the cerebellum retracted. **f,** Soft PEDOT electrode was wrapped around the facial-acoustic nerve complex for subsequent neurophysiological monitoring.

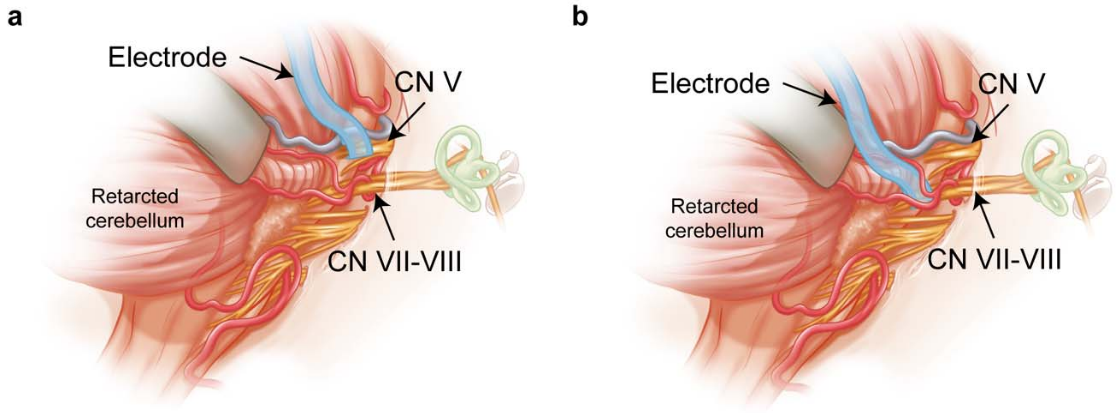


**Supplementary Fig. 5 | Soft PEDOT electrodes for nerve mapping. a-b,** Schematics showing the soft PEDOT electrodes wrapped around the trigeminal nerve (**a**) and the facial-acoustic nerve complex for nerve mapping (**b**).

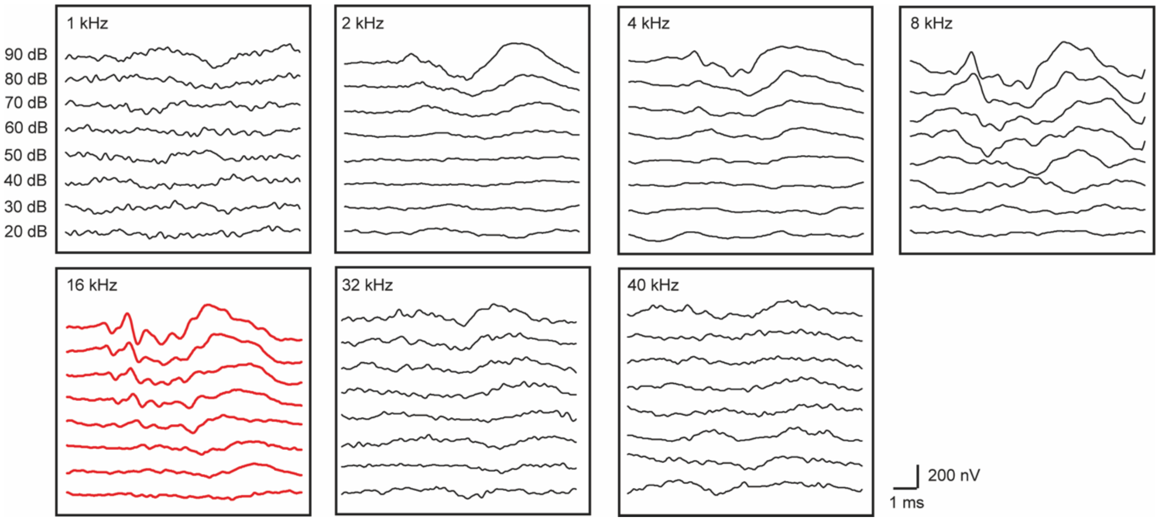


**Supplementary Fig. 6 | Waveforms of BAEP obtained at different frequencies in the rabbit model.** Waveform at 16 kHz was most stable and used in the experiment. BAEP: Brainstem auditory evoked potentials. The data was obtained by conventional rigid electrode.

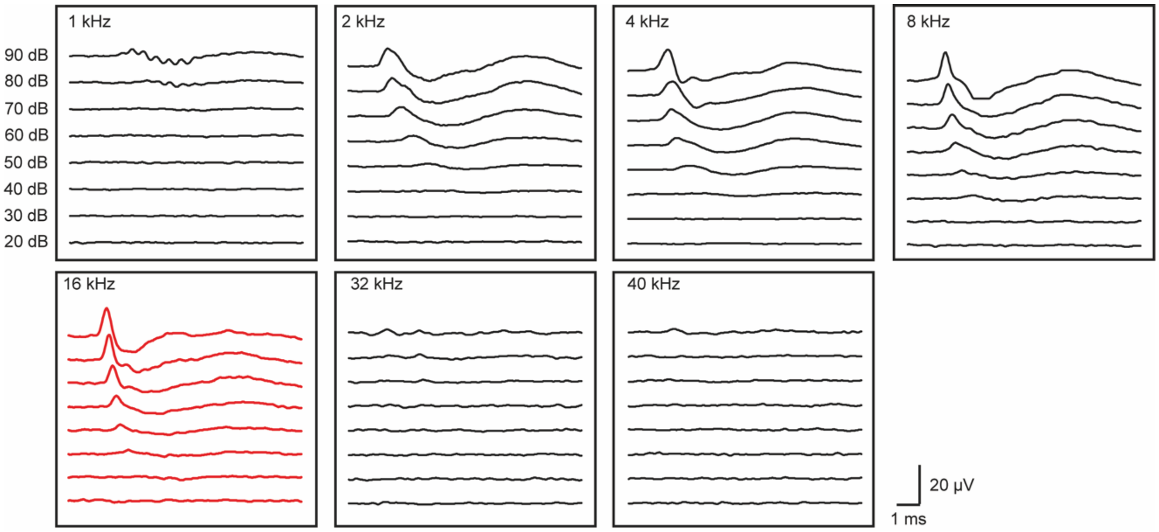


**Supplementary Fig. 7 | Waveforms of CNAP obtained at different frequencies in the rabbit model.** Waveform at 16 kHz was most stable and used in the experiment. CNAP: Cochlear nerve action potentials. The data was obtained by conventional rigid electrode.

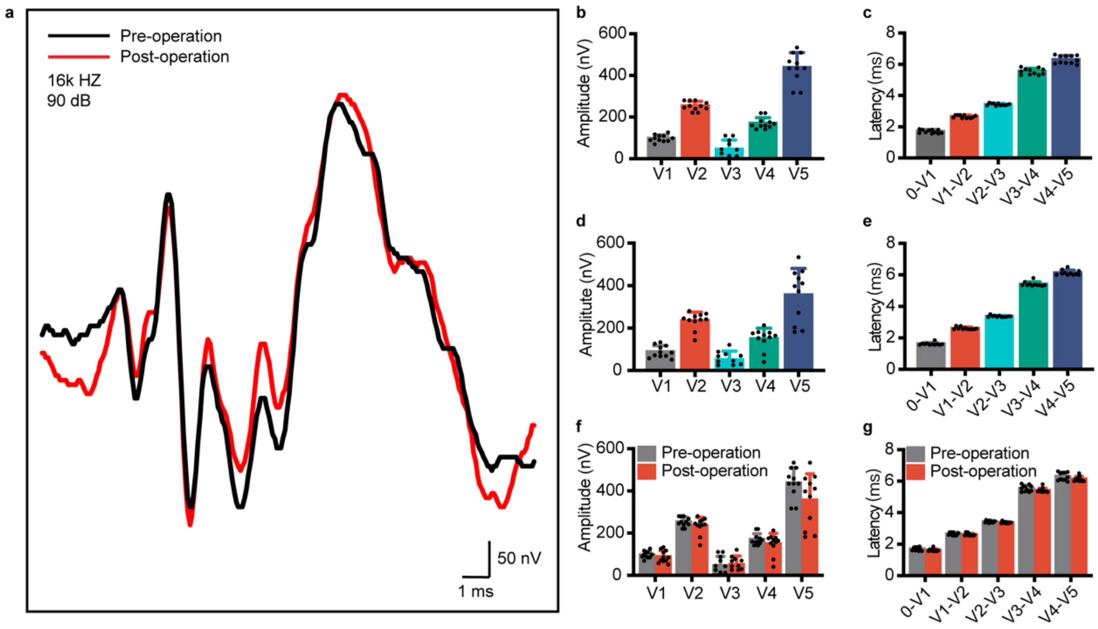


**Supplementary Fig. 8 | Retrosigmoid craniotomy in the rabbit model did not induce hearing loss. a,** BAEP values recorded before and after surgery for comparison. **b-c,** Amplitude (**b**) and latency (**c**) of BAEP before surgery. **d-e,** Amplitude (**d**) and latency (**e**) of BAEP at the time of surgery completion. **f-g,** Comparison of the amplitude (**f**) and latency (**g**) of BAEP before and after surgery. The $P$ values for comparison of the BAEP amplitudes are as follows: V1 ($n = 11$), for pre-operation and the post-operation, $P = 0.353$; V2 ($n = 11$), for pre-operation and the post-operation, $P = 0.202$; V3 ($n = 11$), for pre-operation and the post-operation, $P = 0.790$; V4 ($n = 11$), for pre-operation and the post-operation, $P = 0.234$; V5 ($n = 11$), for pre-operation and the post-operation, $P = 0.076$. The $P$ values for comparison of the BAEP latencies are as follows: 0 - V1 ($n = 11$), for pre-operation and the post-operation, $P = 0.076$; V1 - V2 ($n = 11$), for pre-operation and the post-operation, $P = 0.182$; V2-V3 ($n = 11$), for pre-operation and the post-operation, $P = 0.030$; V3 - V4 ($n = 11$), for pre-operation and the post-operation, $P = 0.092$; V4 - V5 ($n = 11$), for pre-operation and the post-operation, $P = 0.074$. All error bars denote s.d. $*P < 0.05$; $**P < 0.01$; $***P < 0.001$; unpaired, two-tailed Student's t-test was used for f and g. BAEP: Brainstem auditory evoked potentials.

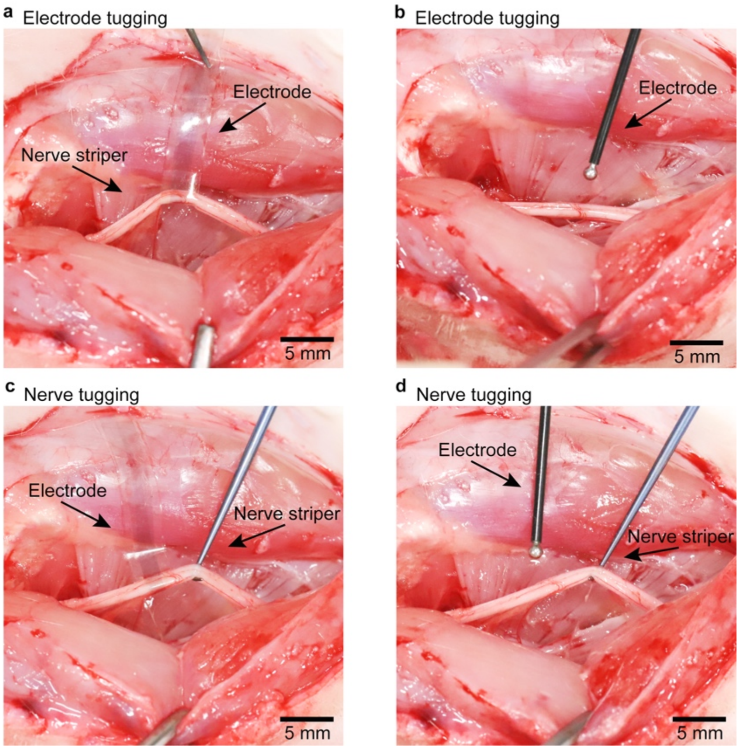


**Supplementary Fig. 9 | Soft PEDOT electrode for robust neural interfaces. a,** Image of soft PEDOT electrode being pulled by tweezer, showing the robustness of the neural interface. **b,** Image of pulling a commercial ball-point electrode, which would cause immediate detachment from the nerve. **c,** Image of a nerve being pulled by a striper, showing the stable contact of the soft PEDOT electrode on the nerve. **d,** Image showing the detachment of a commercial ball-point electrode from a nerve upon pulling by a nerve striper.

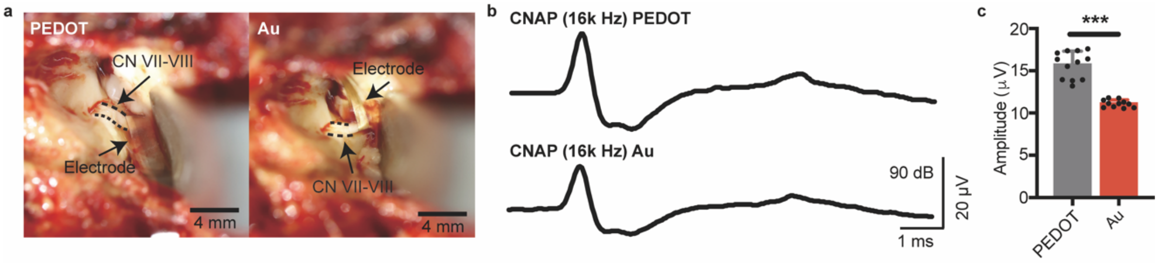


**Supplementary Fig. 10 | Soft PEDOT electrodes for intra-operative CNAP monitoring.** **a,** Photographs of the PEDOT (left) and Au (left) electrode wrapped around the facial-cochlear nerve complex for CNAP recording. **b,** CNAP values were measured from the PEDOT and Au device. **c,** PEDOT electrode could consistently record higher CNAP amplitudes than those from the Au electrode. ($n$ = 11, $P$ < 0.001). All error bars denote s.d. *$P$ < 0.05; **$P$ < 0.01; ***$P$ < 0.001; unpaired, two-tailed Student's t-test was used for **c**. CNAP: Cochlear nerve action potentials.

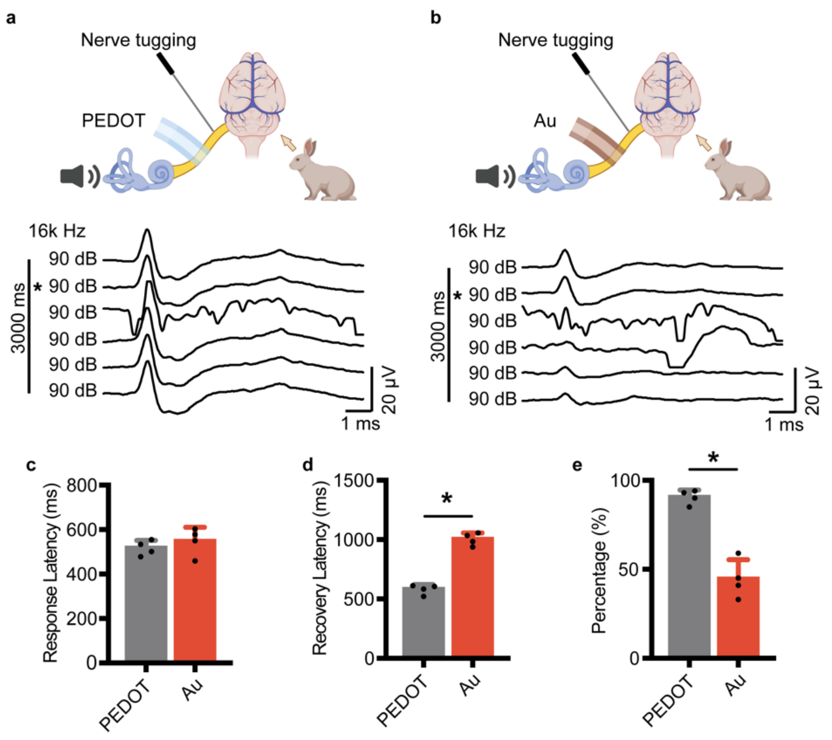


**Supplementary Fig. 11 | PEDOT electrode for real-time auditory monitoring. a,** Schematic and variation of CNAP waveform measured by PEDOT electrode during nerve tugging. * denotes the time point of nerve tugging. **b**, Schematic and variation of CNAP waveform measured by Au during nerve tugging. * denotes the time point of nerve tugging. **c**, Comparison of response latencies of CNAP recorded by PEDOT or Au electrode during nerve tugging. ($n$ = 4, $P$ = 0.486). **d**, Comparison of recovery latencies of CNAP recorded by Au or PEDOT electrode during nerve tugging. ($n$ = 4, $P$ = 0.029). **e**, Percentage of CNAP amplitude after physical nerve tugging recorded by Au or PEDOT electrode. ($n$ = 4, $P$ = 0.029). All error bars denote s.d. *$P$ < 0.05; **$P$ < 0.01; ***$P$ < 0.001; unpaired, two-tailed Student's t-test was used for **c-e**. CNAP: Cochlear nerve action potentials.

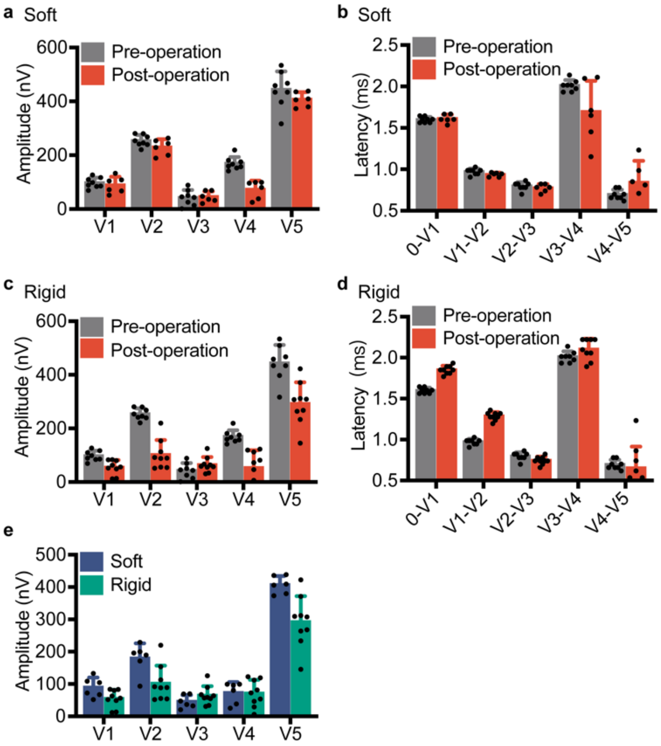


**Supplementary Fig. 12 | Hearing preservation without nerve tissue damage after CINM by soft PEDOT electrode. a-b**, Comparison of BAEP amplitude (**a**) and latency (**b**) with and without soft PEDOT electrode wrapping. **c-d**, Comparison of BAEP amplitude (**c**) and latency (**d**) with and without rigid electrode wrapping. **e**, Comparison of BAEP amplitude with soft PEDOT electrode wrapping and with rigid electrode wrapping. BAEP: Brainstem auditory evoked potentials.

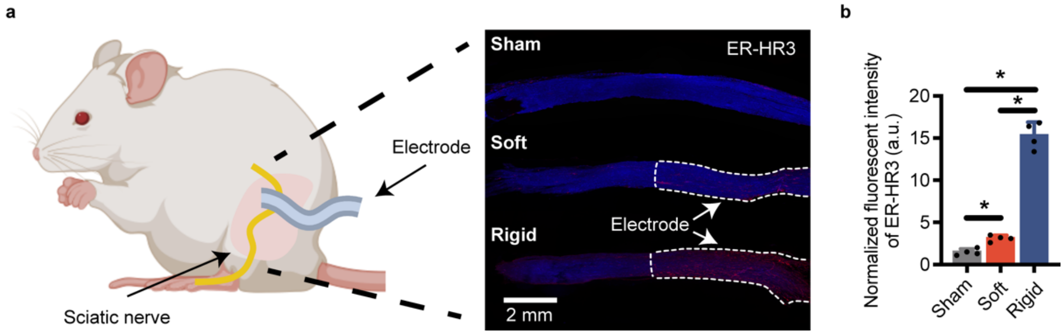


**Supplementary Fig. 13 | Long term biocompatibility. a**, Schematic of the biocompatibility study and longitudinal-section slice of sciatic nerve labeled by the inflammatory biomarker ER-HR3 for soft PEDOT, rigid and sham control. A soft PEDOT electrode or a rigid electrode was wrapped around the sciatic nerve of the mice for 2 weeks, respectively. **b**, Histogram showing the mean fluorescence intensity of ER-HR3 for soft PEDOT, rigid and sham control ($n = 4$). The $P$ values for comparison of the ER-HR3 intensities are as follows: for sham and the soft PEDOT electrode, $P = 0.029$; for sham and the rigid electrode, $P = 0.029$. for soft PEDOT and rigid electrode, $P = 0.029$. All error bars denote s.d. *$P < 0.05$; **$P < 0.01$; ***$P < 0.001$; unpaired, two-tailed Student's t-test was used for **b**.

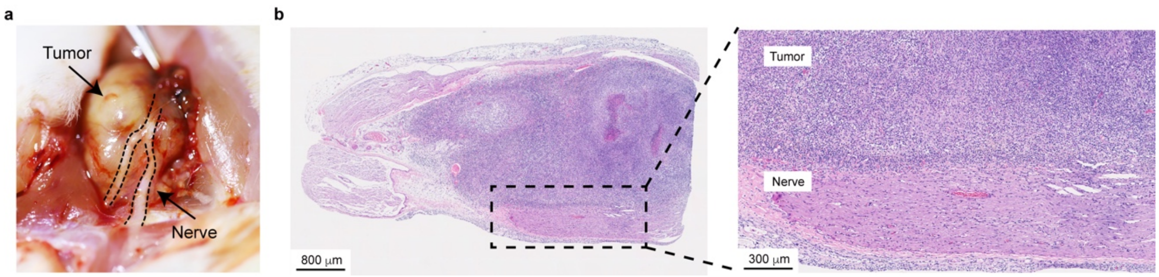


**Supplementary Fig. 14 | Schwannoma on sciatic nerve model. a**, Representative image of Schwannoma animal model. **b**, Cross-sectional slice of tumor bearing sciatic nerve by H&E staining.

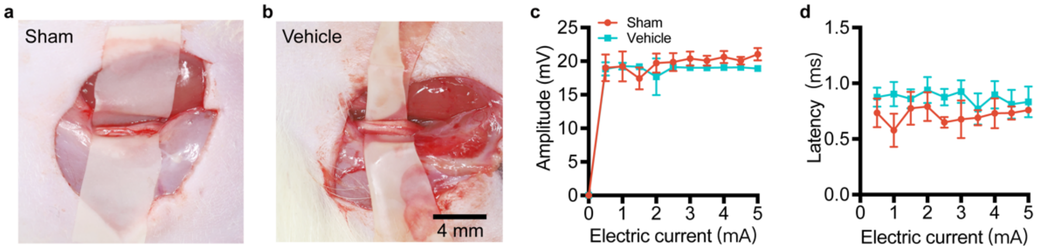


**Supplementary Fig. 15 | EMG study of sciatic nerves between sham and vehicle group.** **a**, Representative image of a normal sciatic nerve. **b**, Representative image of a sciatic nerve 14 days after saline injection. **c**, Comparison of electromyographic amplitude between normal sciatic nerves and sciatic nerves injected with saline at different stimulation strengths ($n$ = 4, $P$ = 0.245). **d**, Comparison of electromyographic latency between normal sciatic nerves and sciatic nerves injected with saline at different stimulation strengths. ($n$ = 4, $P$ = 0.362). All error bars denote s.d. One-way ANOVA with Holm-Sidak's multiple comparisons test was used for **c** and **d**. EMG: Electromyography.

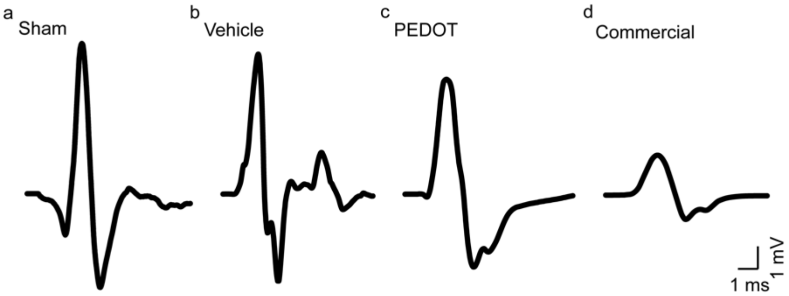


**Supplementary Fig. 16 | Electromyographic waveforms of sciatic nerves in the animal model. a**, Electromyographic waveforms of a normal sciatic nerve (a), a sciatic nerve with saline injection (b), a sciatic nerve tumor model 4 weeks after resection with PEDOT for CINM (c), and a sciatic nerve tumor model 4 weeks after resection with commercial metal electrode for INM (d). In all case, the nerves were stimulated by the same current level of 1.5 mA.

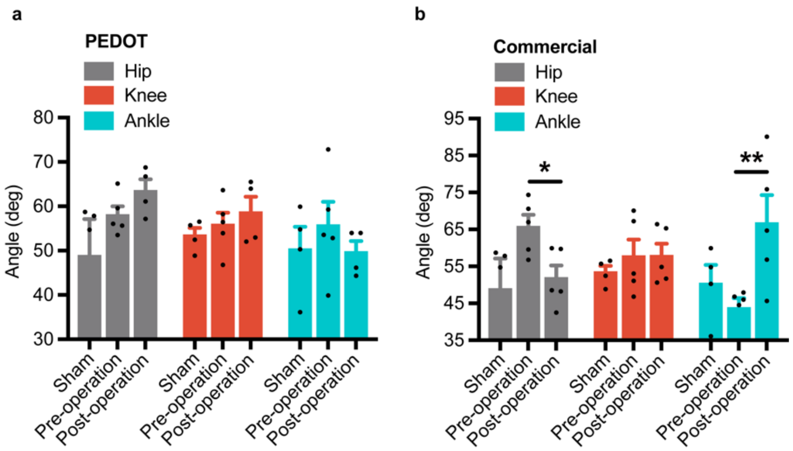


**Supplementary Fig. 17 | Bases for gait recording and data analyses. a**, Average step cycles at the three time points for hip, knee, and ankle joint angles of the mice before and 4 weeks after tumor removal on sciatic nerve by soft PEDOT electrode monitoring compared with sham control. **b**, Average step cycles at the three time points for hip, knee, and ankle joint angles of the mice before and 4 weeks after tumor removal on sciatic nerve by commercial metal electrode monitoring compared with sham control. The *P* values for comparison of the step cycles of hip angles are as follows: for the pre-operation monitored by commercial electrode ($n$ = 5) versus the 4 weeks post-operation monitored by commercial electrode ($n$ = 5), $P$ = 0.020; The *P* values for comparison of the step cycles of ankle angles are as follows: for the pre-operation monitored by commercial electrode ($n$ = 4) versus the 4 weeks post-operation monitored by commercial electrode ($n$ = 5), $P$ = 0.003. All error bars denote s.d. *$P < 0.05$; **$P < 0.01$; ***$P < 0.001$; unpaired, two-tailed Student’s t-test was used for **a** and **b**. The increased hip angle and decreased ankle angle means better gait of mice.

**Supplementary Video 1 | Video showing BAEP monitoring (5× speed playback).**

**Supplementary Video 2 | Video showing CNAP monitoring (0.5× speed playback).**

**Supplementary Video 3 | Gait study of sciatic nerve VS model. Only right leg was injected with tumor cells.**

**Supplementary Video 4 | Gait comparison between rat monitored with PEDOT and commercial metal electrodes at 4 weeks after tumor removal. Only the right leg was injected with tumor cells.**

**Supplementary Video 5 | Electrical stimulations with PEDOT electrode on facial-acoustic nerve complex elicit rabbit whiskers wiggling (amplitude 4 mA, frequency 1 Hz).**

**Supplementary Video 6 | Comparison of electrical stimulation efficiency between soft PEDOT electrode and conventional metal electrode with the same electrode area on the facial-acoustic nerve complex (amplitude 4 mA, frequency 1 Hz). The soft PEDOT electrode can stimulate rabbit whiskers wiggling at a low threshold of 4 mA, while no rabbit whisker wiggling could be observed for the conventional metal electrode.**

### References


1. Correction: Anti–Vascular Endothelial Growth Factor Therapies as a Novel Therapeutic Approach to Treating Neurofibromatosis-Related Tumors. *Cancer Research* **71**, 1197-1197, doi:10.1158/0008-5472.Can-10-4418 (2011).
2. Park, C. *et al.* Protective Effect of Baicalein on Oxidative Stress-induced DNA Damage and Apoptosis in RT4-D6P2T Schwann Cells. *Int J Med Sci* **16**, 8-16, doi:10.7150/ijms.29692 (2019).
3. Akil, O., Oursler, A. E., Fan, K. & Lustig, L. R. Mouse Auditory Brainstem Response Testing. *Bio Protoc* **6**, doi:10.21769/BioProtoc.1768 (2016).
4. Liu, Y. *et al.* Morphing electronics enable neuromodulation in growing tissue. *Nat Biotechnol* **38**, 1031-1036, doi:10.1038/s41587-020-0495-2 (2020).
5. George, P. M. *et al.* Electrical preconditioning of stem cells with a conductive polymer scaffold enhances stroke recovery. *Biomaterials* **142**, 31-40, doi:10.1016/j.biomaterials.2017.07.020 (2017).
6. George, P. M. *et al.* Fabrication and biocompatibility of polypyrrole implants suitable for neural prosthetics. *Biomaterials* **26**, 3511-3519, doi:10.1016/j.biomaterials.2004.09.037 (2005).
7. Gao, X. *et al.* Anti-VEGF treatment improves neurological function and augments radiation response in NF2 schwannoma model. *Proc Natl Acad Sci U S A* **112**, 14676-14681, doi:10.1073/pnas.1512570112 (2015).
8. Wu, L. *et al.* Losartan prevents tumor-induced hearing loss and augments radiation efficacy in NF2 schwannoma rodent models. *Sci Transl Med* **13**, 4816, doi:10.1126/scitranslmed.abd4816 (2021).
9. Akay, T. Long-term measurement of muscle denervation and locomotor behavior in individual wild-type and ALS model mice. *J Neurophysiol* **111**, 694-703, doi:10.1152/jn.00507.2013 (2014).
10. Fiander, M. D., Stifani, N., Nichols, M., Akay, T. & Robertson, G. S. Kinematic gait parameters are highly sensitive measures of motor deficits and spinal cord injury in mice subjected to experimental autoimmune encephalomyelitis. *Behav Brain Res* **317**, 95-108, doi:10.1016/j.bbr.2016.09.034 (2017).
